# Circadian disruption induces sex-specific Alzheimer’s pathophysiology and immune cell reprogramming

**DOI:** 10.64898/2026.04.30.721994

**Authors:** Carmalena V. Cordi, Naomi G. Falkenberg, Gretchen T. Clark, Noah G. Allen, Joshua R. Chuah, Russell Ulbrich, Ava A. Herzog, Meghana Lanka, Emily J. Collins, Marvin Bentley, Jonathan S. Dordick, Meaghan S. Jankowski, Jennifer M. Hurley

## Abstract

Circadian disruption (CD) is increasingly recognized as a sex-specific risk factor for Alzheimer’s disease (AD). However, the mechanisms linking CD to AD, and the role of biological sex in this interaction, are unclear. Immunometabolic regulation is extensively circadianly timed, has sex-specific phenotypes, and plays a role in AD. Therefore, we hypothesized that CD affects the timing of immunometabolism, contributing to the sex-specific effects of CD on AD. To demonstrate this, we subjected male and female APP/PS1 mice to chronic disruptive lighting to model circadian disruption, finding CD induced a female-specific reduction in amyloid plaque burden but an increase in the infiltration of peripheral macrophages into the brain. Concomitantly, we found macrophages exhibited CD-associated immune reprogramming, which in females led to altered immunometabolic timing, an increase of macrophages in the activated state, and elevated levels of reactive oxygen species (ROS), supporting a role for immunometabolism in the sex-specific effects of CD in AD.

**Graphical Abstract:** 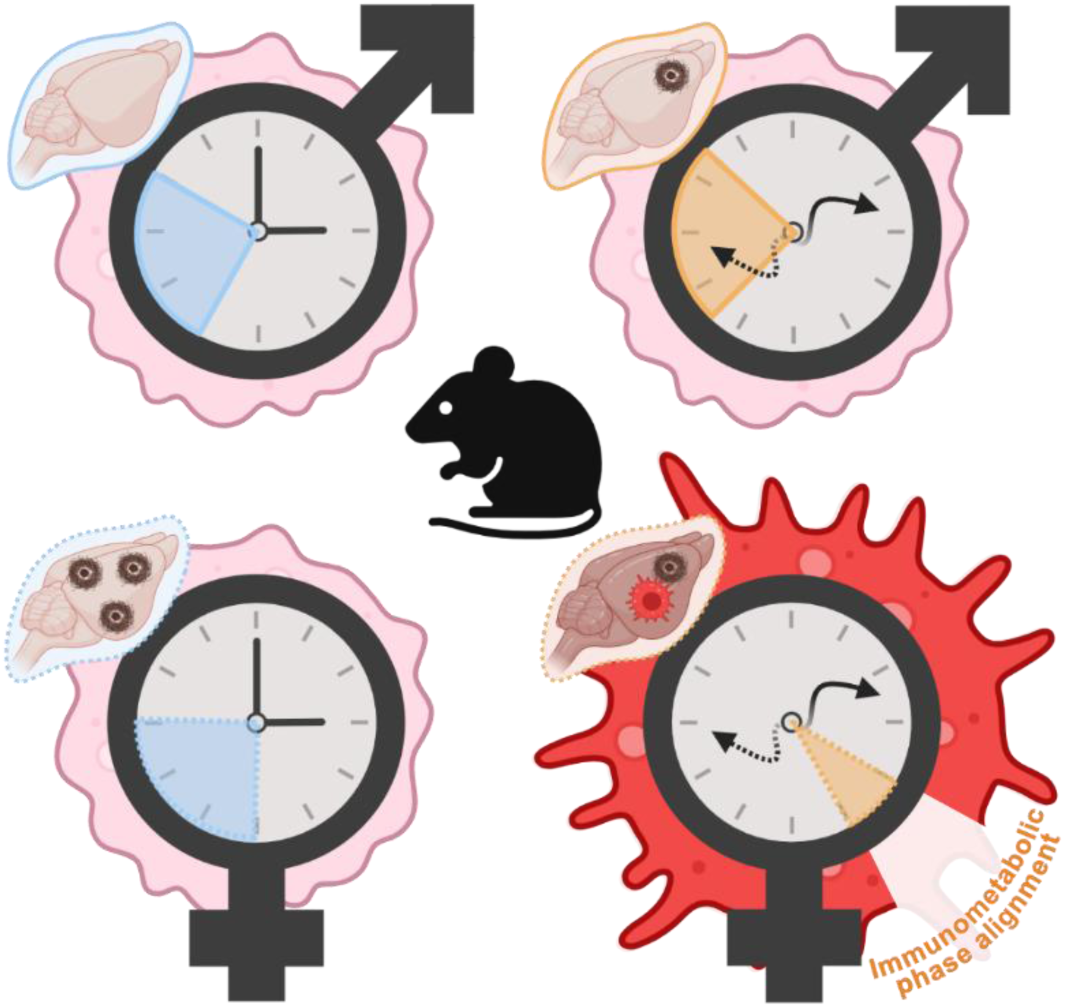

**Highlights:**

- Circadian disruption reduces Aβ plaque load specifically in female mice
- Peripheral immune infiltration correlates with reduced Aβ plaques in females
- Circadian disruption coordinates phasing of circadian immunometabolic proteins
- In females, circadian phase advancement correlates with increased ROS

## Introduction

Circadian rhythms widely regulate metabolic, immune, and neuronal processes across the day-night cycle, tuning physiology to Earth’s daily rhythms.^1^ At the cellular level, these rhythms are generated by the core molecular clock, a transcription-translation feedback loop (TTFL) in which BMAL1 and CLOCK drive rhythmic expression of period (PER1-3) and cryptochrome (CRY1-2) proteins, which in turn inhibit BMAL1/CLOCK activity to generate approximately 24-hour oscillations in transcriptional and post-transcriptional regulation at the cellular level.^2^ Environmental factors such as shift work, irregular light exposure, and chronic sleep-wake disruption result in circadian disruption (CD), which has negative consequences on human health.^3^ CD is associated with systemic inflammation, metabolic dysregulation, and increased risk for neurodegenerative diseases such as Alzheimer’s disease (AD) in aging populations.^4–9^ In fact, individuals chronically exposed to disruptive lighting experience accelerated cognitive decline and elevated AD incidence.^10–13^ However, an understanding of the mechanistic link between CD and AD is limited.

AD is marked by progressive amyloid-β (Aβ) accumulation, tau pathology, and a chronic inflammatory state, highlighting several potential mechanisms that could explain the connection between CD and AD.^7,9,14^ One pertinent mechanism lies in the circadian influence on the immune system, where intrinsic molecular clocks in immune cells drive daily oscillations in the functional immune response, immunometabolic regulation, and inflammation.^15–19^ Importantly, Aβ clearance is dependent on immune cells and has been shown to be clock-regulated.^18^ Circadian disruption alters these rhythms, affecting many aspects of immunity.^20^ Relevant to AD, CD impairs microglial surveillance and cytokine dynamics, and disrupts the activity of both brain-resident and peripheral immune cells.^16,17,21,22^ CD also produces exaggerated inflammatory responses and diminished temporal coordination of innate immunity.^6,23^ These effects are known to compromise neuro- and peripheral-immune homeostasis, which contributes to cognitive deficits in AD.^9,21,24^ Thus, immune dysfunction due to CD is a plausible pathway through which CD may accelerate AD-related processes.

Complicating an understanding of the role of CD in AD are some general sex-specific effects of AD, with female patients with AD exhibiting higher disease incidence, greater severity, and a faster disease progression as compared to their male counterparts.^25–29^ Importantly, there is a female-specific vulnerability to CD, with an increase in the inflammatory response upon CD.^30^ Also, females that regularly undergo shift work have increased rates of AD and other signs of decreased cognition.^31^ In concordance with this, there are known sex-specific differences in the circadian regulation of the immune system, including daily cytokine rhythms and sleep-related inflammatory variation.^32,33^ These effects are exacerbated by the fact that the general female immune response already shows heightened inflammatory reactivity.^34^ However, an understanding of the factors that link CD, sex-specific immune changes, and AD progression in humans is currently lacking.

To investigate this link, we used a disruptive lighting paradigm, similar to what is experienced during shift work, to induce sustained circadian disruption in mice.^35^ Hemizygous APPswe/PS1dE9 (APP/PS1) mice of both sexes (referred to as AP), alongside control littermates (LM), were used to determine the effect of circadian disruption in the preclinical stages of AD. Using this protocol, we integrated longitudinal behavioral circadian profiling with multi-omic and functional analyses of peripheral macrophages, comprehensive immune phenotyping, and histopathological assessment of Aβ plaques and immune cells in the brain to capture the systemic and sex-specific effects of prolonged circadian disruption. Our results reveal that chronic circadian disruption induced sex-specific effects on plaque pathology and immune cell infiltration within the brain, with females displaying lowered amounts of Aβ plaques and increased infiltration of peripheral macrophages upon CD. However, these effects were coordinated with the sex-specific rewiring of circadian regulation in peripheral macrophages, which manifested as altered immunometabolic homeostasis, with females demonstrating an increased phase alignment of immunometabolic pathways and overall activation and inflammatory levels. In total, these results suggest a significant role for peripheral inflammation in the sex-specific effect of CD on AD.

## Results

### Circadian Disruption Sex-Specifically Modulates Aβ Accumulation

To examine the connection between CD, sex-specific immune changes, and AD progression, we first needed a model that would allow us to investigate AD progression. We selected the transgenic hemizygous APPswe/PS1dE9 (APP/PS1) mouse strain, which expresses the human amyloid precursor protein (APP) with the Swedish mutation and presenilin 1 (PSEN1/PS1) with the L166P mutation (JAX MMRRC Stock# 034829).^36^ This APP/PS1 strain, (referred to here as AP), mimics AD phenotypes, alters clock function in later stages of AD, and has higher plaque accumulation/mortality in females.^37,38^ Littermate (LM) controls lacking the APP/PS1 mutations were included for all comparisons. To reduce hormonal variability, group-housed female mice were exposed to male pheromones through soiled bedding to induce the Whitten effect, which allowed for the synchronization of the female estrus cycle during sample collection to just after peak estrogen levels, minimizing hormone variability (**Figure 1A**).^39,40^

**Figure 1.**
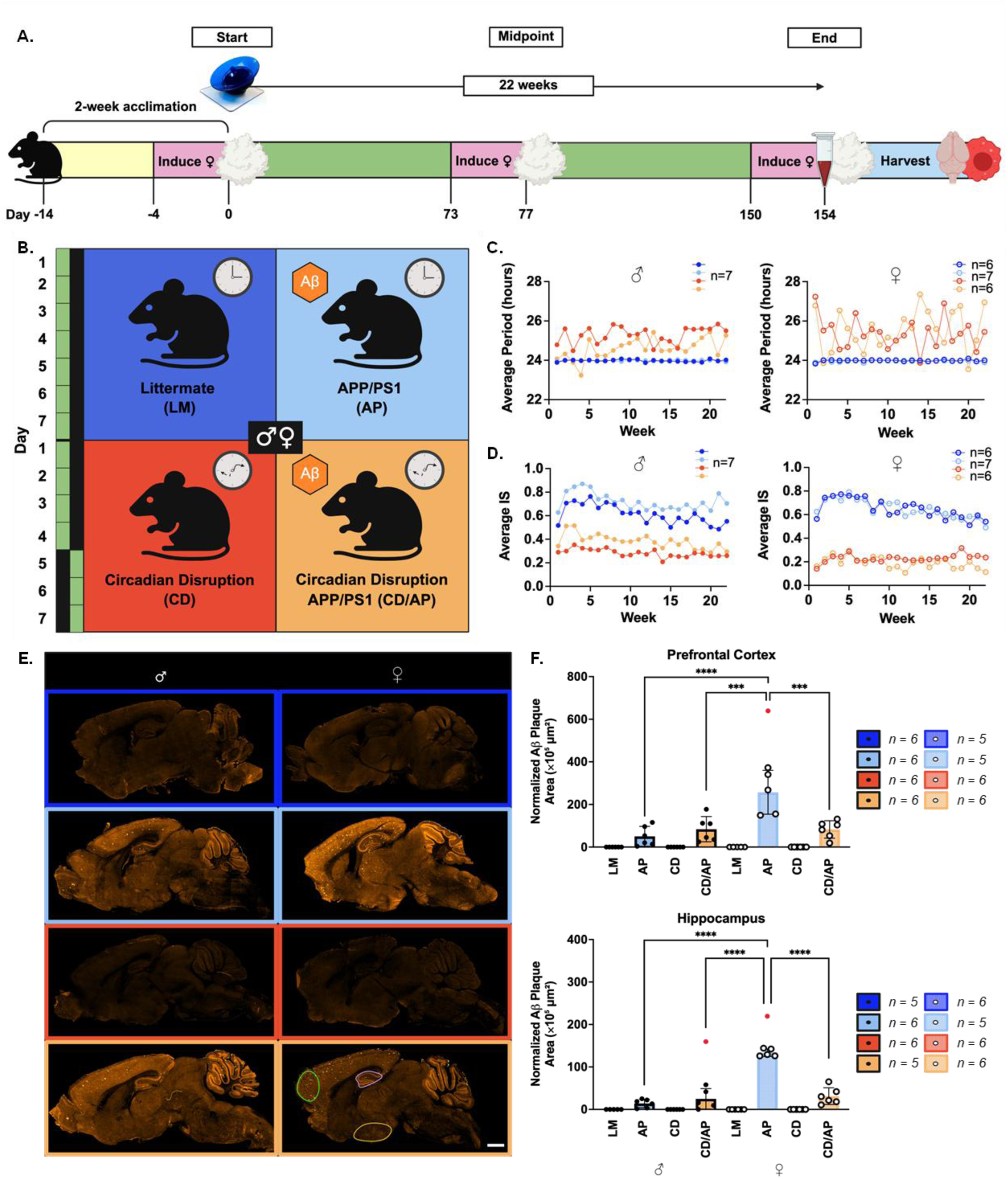
Chronic circadian disruption disturbs circadian rhythms and sex-specifically alters Aβ plaque accumulation. (A) Schematic of lighting protocol and experimental procedures. Cotton represents male soiled bedding, wheel represents monitoring of activity, and organs represent tissue sampling. (B) A diagram depicting the setup for the treatment groups and lighting protocols, with 12-hour increments of “lights on” (green) and “lights off” (black) for each day. (C) Mean weekly periods and (D) Interdaily Stability (IS) assessed over 22 weeks (*n* = 7 except female LM and CD/AP where *n* = 6). (E) Representative 25 μm sagittal images of immunolabeled brain sections (scale bar = 1 mm), with regions indicated: prefrontal cortex (green), hippocampus (purple), and hypothalamus (yellow). (F) Total Aβ plaque load in the prefrontal cortex and hippocampus normalized to the region of interest (*n* = 6 except female AP prefrontal cortex and male LM hippocampus where *n* = 5). * = p < 0.05, ** = p < 0.01, *** = p < 0.001, and **** = p < 0.0001. Red dots represent outliers. Bars show mean ± SD. Dark blue/black outline = male LM, light blue/black outline = male AP, red/black outline = male CD, orange/black outline = male CD/AP, dark blue = female LM, light blue = female AP, red = female CD, orange = female CD/AP. ♂ = male, ♀ = female. See also Figure S1 and Table S1.

To impose CD on murine physiology in a manner mimicking the effect of human-built lighting on humans, mice were group-housed by sex and genotype in standard cages positioned between diffused monochromatic green LED lights (**Figure S1**) (525 nm, 4 μW/cm²).^35^ Our lighting protocol consisted of 22 weeks of a standard 12 hours of light and 12 hours of dark (12L:12D) schedule for seven days for the control population, while the CD population experienced 22 weeks of a rotating pattern of four days of 12L:12D followed by three days of 12D:12L (**Figures 1A** and **1B**). This lighting protocol was formulated based on three factors. First, preliminary data demonstrated a rotating green-light cycle consisting of four days of 12L:12D followed by three days of 12D:12L led to elevated inflammatory serum cytokines (e.g., CXCL10 and IL-15) (**Figure S1**).^41–43^ A similar trend has also been reported in humans.^44^ Second, APP/PS1 mice exhibit Aβ deposition (including in the suprachiasmatic nucleus (SCN)), synaptic loss, and gliosis beginning between 6-9 months of age.^36^ Finally, the age of the mice in our study (∼8 months) means that they are just before the mouse endocrine equivalent of human perimenopause (roughly 9 months of age).^45,46^ Therefore, our protocol length allowed for the evaluation of the effect of CD on Aβ accumulation without the confounding feed-forward effect of Aβ accumulation in the SCN or decreasing estrogen levels.

To confirm the effectiveness of our CD protocol in disrupting circadian rhythms, we monitored the wheel revolutions for a subset of mice with running wheels in ten-minute intervals over 22-weeks. Using this wheel running data, we conducted periodogram analyses to determine period length and interdaily stability (IS) (the consistency of activity between days, with higher IS suggesting a stronger circadian consistency) (**Figures 1A**, **1C**, and **1D**).^47^ As expected, both male and female mice in the 12L:12D lighting protocol (LM and AP) maintained consistent 24-hour rhythms throughout the 22 weeks, while the mice exposed to the disruptive lighting protocol (CD and CD/AP) showed more variable periods (23-28 hours) (**Figures 1C** and **1D**). IS generally declined over time in all groups, consistent with known aging-related circadian effects,^48^ but remained lower under disruptive lighting conditions (CD and CD/AP) compared to the standard lighting (LM and AP), indicating that CD disrupted day-to-day activity patterns (**Figures 1C** and **1D**). These data demonstrated that our green-light CD lighting protocols successfully disrupted circadian rhythms in both the AP and LM mice.

To examine the effect of CD on gross cognition, we subjected our non-wheel running mice to a nesting test at the beginning, midpoint, and end of the 22-week lighting protocol (**Figure S1 C** and **D**). Each mouse was temporarily individually housed with four pre-weighed squares of nestlet material for 24 hours. We then scored the nests blindly for construction quality on a scale from one to five and weighed the remaining nestlet material to determine the amount used.^49^ We found no significant differences in nestlet scores in all sexes, genotypes, and lighting conditions over time, indicating that gross cognitive function remained intact in our early AD model, regardless of the lighting schedule, aligning with the known data on this model ^50^. This is bolstered by the consistent period and IS score that is seen in the aged (22-week-old) mice (**Figure 1C** and **1D**). In total, our protocol allowed us to analyze the effects of CD in an AP context prior to the confounding effects of AD on the functionality of the circadian clock.

Given the lack of gross cognitive effect, we next examined if there was a pathophysiological change in the mice that had been subjected to CD. As Aβ buildup is considered one of the earliest signs of AD,^51^ we examined differences in Aβ plaque accumulation in the brain. To do so, after our 22-week lighting protocol, we sacrificed the mice and dissected whole brain tissue, which was immediately fixed with paraformaldehyde and sucrose. The fixed brains were frozen in OCT, bisected, and sectioned along the sagittal plane, allowing for the examination of the prefrontal cortex, hippocampus, and hypothalamus, all within the same section. Each brain section was then labeled with a 6E10 antibody to detect Aβ plaques *in situ* (**Figure 1E**). We then imaged using Tissue Gnostics TissueFAXS and analyzed three distinct regions, the prefrontal cortex, hippocampus, and hypothalamus, with Tissue Gnostics StrataQuest used to identify dense peptide areas indicative of Aβ plaque load (**Figures 1E**, **1F** and **S1F**). These regions were chosen as the prefrontal cortex is essential for cognitive functions, the hippocampus is critical for memory and learning, and the hypothalamus regulates homeostasis and circadian rhythms via the SCN.^52,53^ Further, these areas are sequentially impacted by Aβ accumulation during the progression of AD, with Aβ accumulating first in the prefrontal cortex, then the hippocampus, and finally in the hypothalamus.^54–56^ Consequently, our analysis of these regions enabled us to assess the state of Aβ buildup in early AD model mice.

As anticipated because Aβ does not naturally accumulate in mice, we noted no Aβ plaques in the LM mice, male or female (**Figure 1F**). Similarly, as accumulation of Aβ plaques in the hypothalamus occurs only after nine months, the minimal accumulation of Aβ plaques in the hypothalamus (both males and females) we found was expected (**Figure S1 F**).^54–56^ In the prefrontal cortex and hippocampus of AP male mice, we found a measurable Aβ plaque load in the mice that underwent the standard 12L:12D lighting protocol but noted an increase in Aβ plaque load when these male mice underwent CD, though this increase was not significant (by Tukey’s multiple comparisons test) (**Figure 1F** and **Table S1**). This suggests in males that CD leads to an increase in Aβ plaque load as early as ∼8 months, in parallel with what has been previously noted ^57,58^.

However, Aβ plaque load in the female mice followed a distinctly different trend. We noted a significant increase in Aβ plaques in the females that received the standard 12L:12D lighting protocol as compared to the male Aβ plaque load in either the AP or AP/CD conditions (by Tukey’s multiple comparisons test). In fact, the total plaque load was three and a half times higher in the prefrontal cortex and five times higher in the hippocampus compared to male mice, regardless of the lighting condition of the males (**Figure 1F**). This result parallels the finding that AD in females has an earlier onset and worse phenotype.^59,60^ When we next tracked Aβ plaque load in the female AP mice exposed to CD, we were surprised to see a decrease in Aβ plaque load as compared to the AP females, with plaque loads in the AP/CD female mice paralleling the range seen in the male AP mice (**Figure 1F** and **Table S1**). These differences were supported by an ANOVA test, which revealed significant interactions among sex, genotype, and lighting in Aβ plaque load in the prefrontal cortex and hippocampus (**Table S1**). This also countered our prediction that CD would increase Aβ plaque load, based on the known worsened AD endophenotypes in females exposed to CD.^31^

### CD and AP Lead to Increased Peripheral Macrophage Infiltration in the Female Mouse Brain

Given the well-documented correlation between negative endophenotypes in AD and Aβ plaque loads, our finding of decreased Aβ plaques in the CD/AP females was incongruous with the demonstration that CD has a negative effect on AD endophenotypes in women.^31,61^ To explore potential mechanisms underlying this discrepancy, we first tested whether an increased microglial presence could have promoted Aβ plaque clearance. To investigate this, we created sagittal brain sections and performed immunofluorescence analysis to count the total microglia (TMEM119^+^/IBA1^+^) (**Figure 2A**) within the same three brain regions, the prefrontal cortex, hippocampus, and hypothalamus (**Figure 2B**). A three-way ANOVA revealed no significant interaction between genotype, sex, and lighting on microglial abundance, with only a modest difference observed between AP and CD/AP females in the prefrontal cortex and none in the hippocampus or hypothalamus (**Figure 2C** and **Table S2**), suggesting that microglia were not the primary drivers of the decreased Aβ plaque load in CD/AP females.

**Figure 2.**
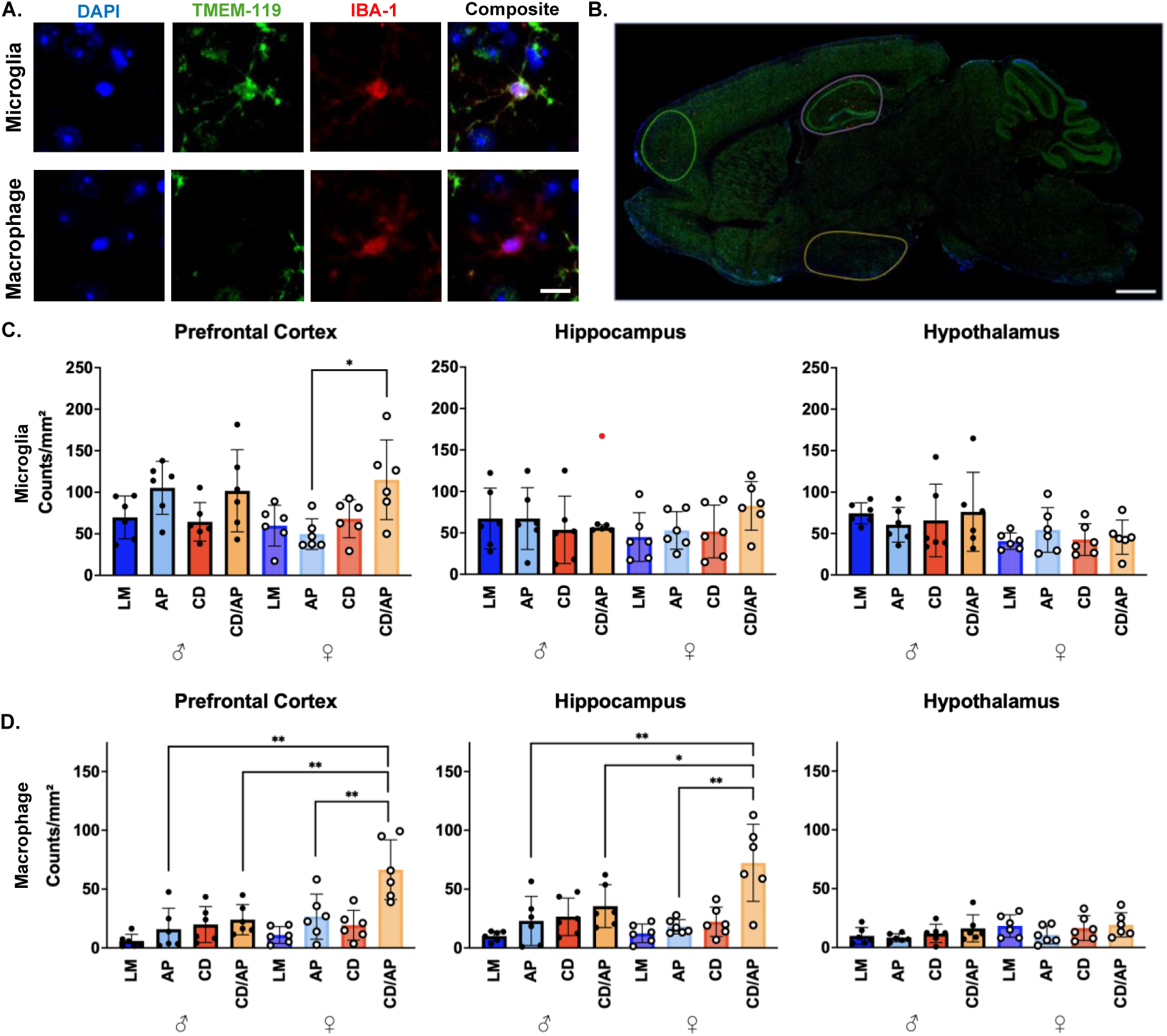
CD in an AD context in females leads to increased infiltration of peripheral macrophages into the brain. (A) Representative TMEM119, IBA1, and DAPI nuclear labeled images of individual cells counted as either microglia (top) or macrophages (bottom) (scale bar = 10 μm). Individual channels in (A) are shown along with a composite image of all three labels (white scale bar = 10 μm). (B) A composite image of a 25 μm sagittal brain (scale bar = 20 µm) with the brain regions labeled: prefrontal cortex (green), hippocampus (purple), and hypothalamus (yellow). (C) Microglia and (D) peripheral macrophage counts normalized to the region of interest (*n* = 6) (* = p < 0.05 and ** = p < 0.01). Red dots represent Dixon Q outliers. Bars show mean ± SD: Dark blue/black outline = male LM, light blue/black outline = male AP, red/black outline = male CD, orange/black outline = male CD/AP, dark blue = female LM, light blue = female AP, red = female CD, orange = female CD/AP. ♂ = male, ♀ = female. See also Figure S2 and Tables S2 and S3.

With the microglia unlikely to be responsible for the difference between the Aβ plaque load in the AP and CD/AP females, we looked for other potential sources for this discrepancy. Recent evidence has shown that peripheral macrophages can migrate to the brain and contribute to Aβ clearance, with peripheral macrophages clearing Aβ more efficiently than microglia.^62,63^ Our data also demonstrated an increase in cytokine levels that signal peripheral immune recruitment (**Figure S1C**). We therefore next used subtractive analysis on immunofluorescently labeled brains to distinguish microglia (TMEM119^+^/IBA1^+^) from infiltrating macrophages (TMEM119^-^/IBA1^+^) (**Figure 2A**), with antibody specificity confirmed using bone marrow-derived macrophages (BMDMs) (**Figure S2**). A three-way ANOVA revealed a significant interaction between genotype, sex, and lighting in macrophage abundance in the prefrontal cortex and hippocampus (**Table S3**), and post-hoc Tukey’s tests showed significantly higher macrophage counts in CD/AP females in these regions compared to all other groups (**Figure 2D**). No differences were observed in the hypothalamus, consistent with the lack of Aβ plaque differences in the area in early AD. In males, peripheral macrophage levels mirrored the similar Aβ plaque loads between AP and CD/AP. Together, these findings indicate that peripheral macrophages infiltrate the brain in significantly higher numbers in female CD/AP mice, which may contribute to the reduced Aβ plaque load in this group.

### CD and AP Affect Circadian Output in Peripheral Macrophages

The incongruity of a reduction in Aβ plaques in CD/AP females with the known negative effects of CD on AD endophenotypes in women drove us to investigate the molecular and behavioral characteristics of the invading peripheral macrophages that may lead to these negative endophenotypes.^31,61^ To do so, bone marrow was harvested from animals in each condition at endpoint, differentiated into BMDMs, and circadianly synchronized via serum shock.^18,19^ The resultant BMDMs were collected every 3 hours over 24 hours and analyzed by Tandem Mass Tag (TMT) mass spectrometry. TMT-based quantitative proteomic data were processed with Learning and Imputation for Mass-spec Bias Reduction (LIMBR) to reduce batch effects,^64^ and each protein’s rhythmicity was assessed using the Extended Circadian Harmonic Oscillator (ECHO) model (**Figure 3A**).^65^

**Figure 3.**
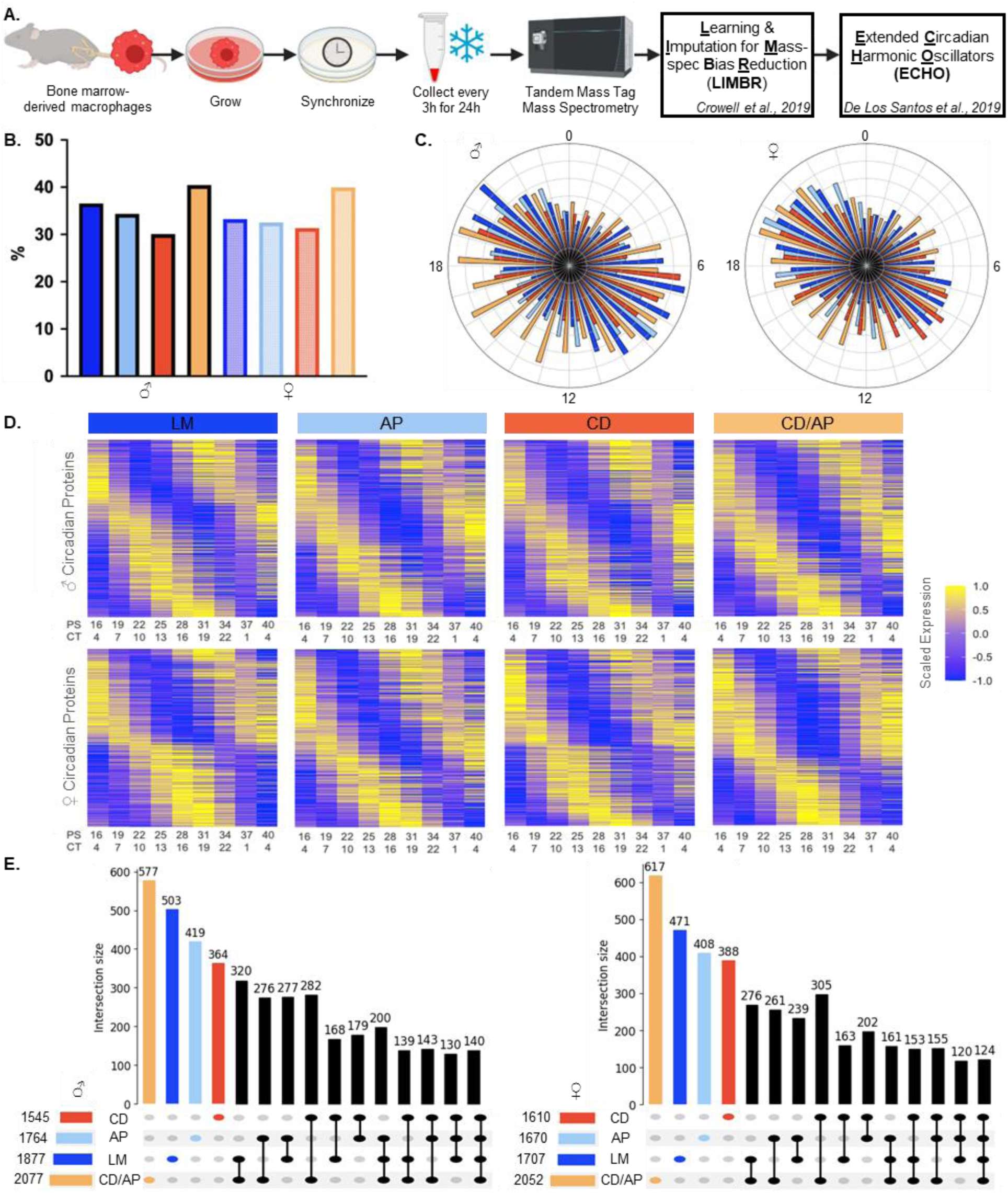
Total proteomics analysis of macrophages over time demonstrates sex, CD, and AP effects on circadian regulation. (A) A schematic representing the workflow of the TMT analysis of BMDMs. (B) The total number of circadian proteins (harmonic/damped/forced) in each of the categories, represented in a bar chart. Dark blue/black outline = male littermate (LM), light blue/black outline = male APP/PS1 (AP) mice, red/black outline = male circadianly disrupted (CD), orange/black outline = male CD/AP, dark blue = female littermate (LM), light blue = female AP, red = female CD, orange = female CD/AP. (C) Radial histograms of the number of oscillating proteins that reach their peak phase at a given circadian timepoint (CT). (D) Heatmaps of proteins with significant circadian oscillations. Scaled expression values (-1 to +1) are shown for post-serum-shock and CT time points. Proteins are ordered along the y-axis by peak phase timing. (E) UpSet plot displaying the number of proteins that are rhythmic across each group, separated by males and females. Fill color indicates genotype/lighting condition for C-E (dark blue = LM, light blue = AP, red = CD, orange = CD/AP). ♂ = male, ♀ = female. See also Figures S3 and S4 and Tables S4 and S5.

To understand how the core molecular clock of a macrophage responded to the combined effects of sex, disruptive lighting, and/or amyloid pathology, we examined circadian oscillations of the key TTFL proteins (BMAL1, CLOCK, CRY1, CRY2, PER1, and PER3) in the BMDMs. As TTFL proteins are expressed in relatively low abundance and were inconsistently detected across conditions, LIMBR-based imputation was not applied for this analysis. Instead, oscillatory behavior was assessed directly from normalized TMT reporter intensities using ECHO.^65^ In general, we found that male and female macrophages preserved oscillations in the proteins within the TTFL, though there was some condition-specific remodeling in the case of CRY2 during CD **(Figure S3** and **Table S4)**.

Having noted few differences in the core clock proteins, we next examined the circadian regulation of the total macrophage proteome from the male and female LM, AP, CD, and CD/AP conditions after LIMBR batch-effect removal.^64^ Based on the observed periods of core clock proteins, we defined a circadian oscillation as a period between 18 and 26 hours (**Figure S3** and **Table S4**). We then used ECHO to model these oscillations (damped, harmonic, or forced, BH adj. p-value <0.05) (**Table S5**).^65^ While the exclusion of overexpressed and repressed proteins slightly reduced the number of identified rhythmic proteins (**Table S5**), it maintained high confidence in rhythmic assignments without affecting downstream ontological analyses.

Using this approach, we reliably detected (in >70% of samples across all datasets) the same 5,136 proteins, with between 1,545 to 2,077 circadian proteins oscillating in any given dataset. We found the CD/AP groups (both male and female) consistently exhibited the highest number of oscillating proteins (**Figure 3B** and **Table S5**). First, to examine differences in the global timing of circadian proteins, we analyzed the timing of peak relative abundance for all oscillating proteins (**Figures 3C** and **3D**). Peak phases were generally bimodal across conditions, occurring around CT9 and CT21. The biphasic nature was similar to our previous analysis in young males (3-6 months), but there was a distinct difference in phase, suggesting that age has an overall effect on circadian phasing in males.^19^ In both male and female CD/AP conditions, the peak of protein phases were broadened between CT6 and 18. Overall, these findings indicate that the global temporal phasing regulation of the proteome is largely conserved across conditions, aligning with the lack of change noted in the oscillation of the TTFL proteins.

Next, to investigate the changes in the proteins that fell under circadian control uniquely within each sex and condition, we generated UpSet plots, highlighting uniquely rhythmic proteins in males and females (**Figure 3E**). In both sexes, most oscillating proteins were unique to a single group, though there was extensive overlap between each of the conditions. To focus on these differences, we employed over-representation analysis of these unique proteins. This analysis demonstrated limited enrichment but did reveal some significant sex and condition-specific changes related to cell cycle/mitosis and immune regulation (BH-adjusted *p* < 0.05; **Figure S4**). These results indicate that while there are significant changes in the rhythmic proteins from each sex and condition, these uniquely circadian rhythmic proteins do not clearly point to a mechanism to explain the incongruity of reduced Aβ plaques in CD/AP females and increased peripheral macrophages.

### CD Drives coordinated Phase Advances in the Female AP Genotype

With our finding of limited effect from either the global protein phasing or uniquely rhythmic proteins, we next examined the circadian proteins shared across standard and disruptive lighting conditions to investigate whether altered phasing of the shared rhythmic proteins could alter physiology in a way that could explain this incongruity. To do so, peak phase (in CT) was plotted for each protein that was common across the experimental (CD) and control conditions, and then heatmaps were used to visualize concentrated regions of phasing (**Figure 4A**). Concentration of points along the line of identity represent little to no change in phase, while points off the line of identity represented changes to the phase. In LM versus CD comparisons, both males and females exhibited a strong peak on the line of identity between CT 12–21 with a minor secondary peak on the line of identity between CT 3-6, suggesting little phase shift in the shared circadian proteins. In AP versus CD/AP, males exhibited a strong peak on the line of identity between CT 3-6 with a secondary peak on the line of identity between CT 15-18, again suggesting little phase shift in the shared circadian proteins. However, while the AP versus CD/AP female exhibited a strong peak on the line of identity between CT 3-6 and a secondary peak on the line of identity between CT 15-18, they also exhibited a peak off the line of identity in the upper left-hand quadrant, suggesting a group of proteins undergoing phase advancement (**Figure 4A**).

**Figure 4.**
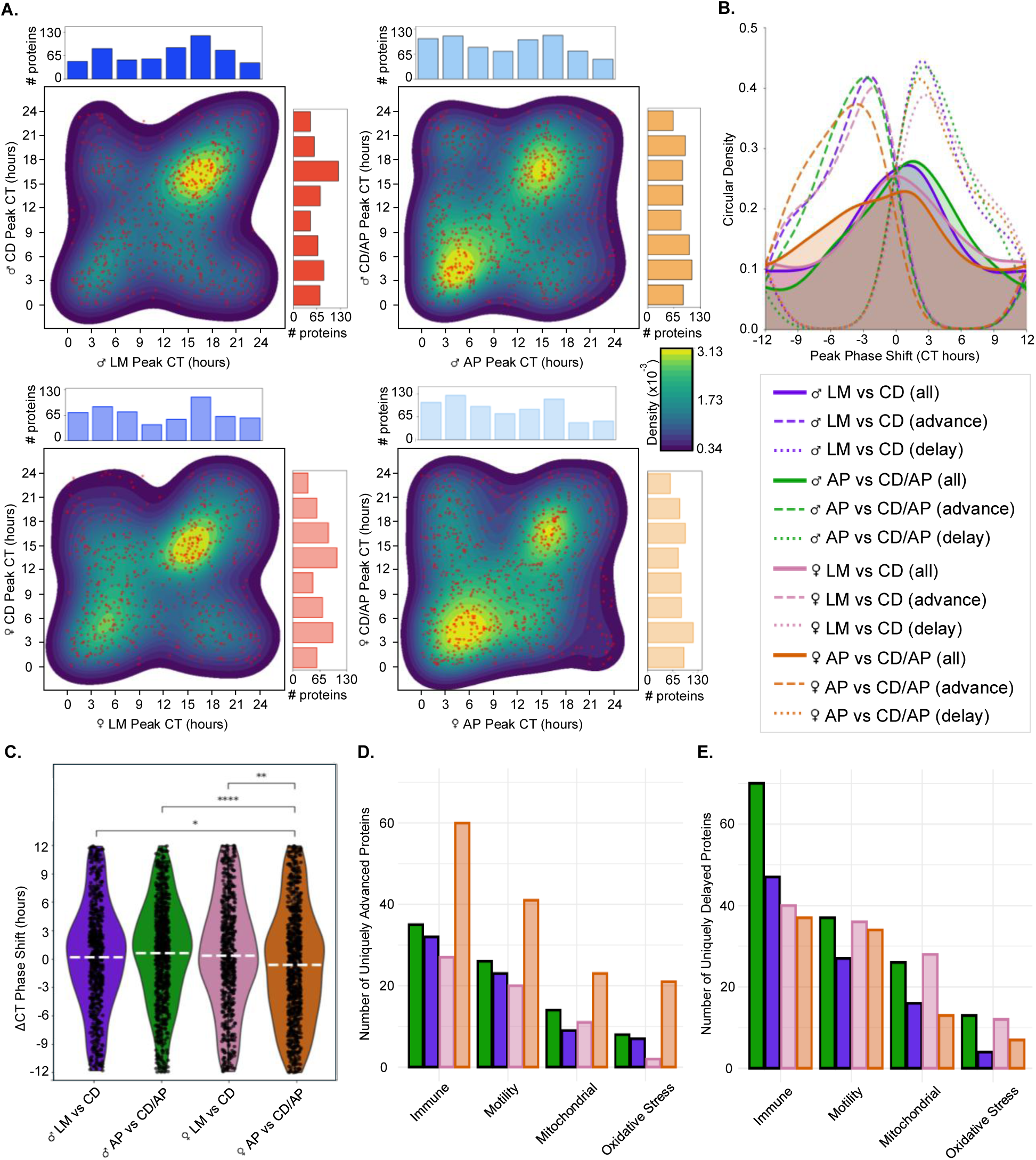
CD sex-specifically affects circadian protein phase timing in AP. (A) Density scatter plots and marginal histograms of the phase timing of the shared circadian proteins compared between male LM vs CD, male AP vs CD/AP, female LM vs CD, and female AP vs CD/AP, with control on the x-axis and CD on the y-axis. The underlying heatmaps depict binned peak phase values by CT, with blue-to-yellow gradients representing low-to-high counts. Marginal histograms show peak phase protein counts in 3-hour CT bins. Dark blue = LM, light blue = AP, red = CD, orange = CD/AP. (B) A density graph of the ΔCTs for the circadian protein comparisons for all (solid lines), phase advanced (ΔCT < 0 h, dashed lines) or phase-delayed (ΔCT > 0 h, dotted lines) proteins. All ΔCTs were mapped onto a normalized 12-h linear scale representing the 24 h circadian cycle. (C) Violin plots of the distribution of the ΔCTs for the circadian protein comparisons, with dashed white lines representing the mean. Individual protein values are displayed as jittered points to illustrate within-group variability. Significant pairwise comparisons are annotated, with * = p < 0.05, ** = p < 0.01, *** = p < 0.001, **** = p < 0.0001 (Dunn’s post-hoc test FDR-adjusted). (D) Bar plots depict the number of unique proteins classified as phase-advanced (ΔCT < −3 h) or (E) phase-delayed (ΔCT > +3 h) for the circadian protein comparisons across four functional groups: Immune, Motility, Mitochondrial, and ROS/oxidative stress. Bar heights represent the number of unique genes assigned to each functional group per phase shift category and comparison. Color indicates lighting comparison purple = ♂ LM vs CD, green = ♂ AP vs CD/AP, pink = ♀ LM vs CD, orange = ♀ AP vs CD/AP). ♂ = male, ♀ = female. See also Table S6.

To quantify these phase shifts, we calculated a ΔCT for all proteins from the above comparisons, whereby phase-advanced proteins (a negative ΔCT) peaked earlier under disruptive lighting and phase-delayed proteins (a positive ΔCT) peaked later (**Figure 4B** -solid lines). We found that in the male and female LM comparisons that phase-advanced and phase-delayed proteins were balanced, as was the case in the male AP versus CD/AP group. However, the female AP versus CD/AP proteins exhibited a group of proteins with a pronounced phase advance (**Figure 4B**). This data was further evident when we plotted the phase delayed or advanced proteins separately on the graph (**Figure 4B** – dashed=advanced, dotted=delayed). The mean ΔCT value of the comparisons further reflected this, as the mean ΔCT from the female AP versus CD/AP was μ = −1.38 h while all other mean ΔCTs were positive (circular standard deviations (σ) and coherence (ρ; mean vector length from circular data) are as follows: ♂ LM vs CD are: μ = 0.36 h, σ = 5.86 h, ρ = 0.309; ♂ AP vs CD/AP: μ = 1.16 h, σ = 5.54 h, ρ = 0.349; ♀ LM vs CD: μ = 0.70 h, σ = 6.36 h, ρ = 0.250; ♀ AP vs CD/AP: μ = −1.38 h, σ = 6.60 h, ρ = 0.224). However, the circular standard deviations indicated broad distributions of phase shifts and low coherence values (**Figure 4B**), revealing heterogeneous timing across proteins. Together, these results demonstrated that although many proteins retain similar phasing between lighting conditions, CD led to a large group of phase-advanced proteins specifically in the AP females.

To further quantify these differences, we examined the distributions of ΔCT phase-shift values using violin plots (**Figure 4C**). The phase-shift values violated assumptions of normality (Shapiro–Wilk p < 0.001) and homogeneity of variance (Levene’s test p < 0.01), so we assessed the between group differences with a Kruskal–Wallis test (**Table S6**). The omnibus test indicated significant differences across the four groups (*H*(3) = 21.31, p *=* 9.1 × 10^-5^). Dunn’s post-hoc tests with FDR correction revealed that the female AP vs CD/AP pair differed significantly from each of the other three pairs (all FDR-adjusted p < 0.014), while no other pairwise comparisons were significant (all FDR-adjusted p > 0.21). These results indicated an AP female-specific circadian phase advance under disruptive lighting conditions.

Next, we evaluated whether the shared phase-shifted proteins we identified had coordinated functional groupings. Using Gene Ontology (GO) Biological Process annotations from *Mus musculus* (org.Mm.eg.db),^66–68^ proteins that underwent a phase advance (ΔCT < −3 h) or phase-delay (ΔCT > +3 h) from all four comparisons were assigned to four broad functional categories: immune activity, motility/cytoskeleton, mitochondrial function, and oxidative stress (**Figures 4D and 4E)**. Among proteins with a phase advance under CD, the female AP versus CD/AP comparison displayed the largest number of early-peaking proteins across these four functional groups, whereas the other pairings (female LM versus CD, male LM versus CD, male AP versus CD/AP) showed more modest changes **(Figure 4D)**. Beyond an increase in immune proteins in the CD/AP males, no analogous trend was observed for the phase-delayed proteins **(Figure 4E)**, indicating a specific phase advance of proteins in the AP females across immune and metabolic pathways due to CD.

### Pathway-Level Circadian Protein Ontological Timing Demonstrates Female-Specific Phase Coordination

Given the immune and metabolic specific changes that we noted in the phase-advanced female CD/AP circadian proteins, we wondered if there was a phase-specific enrichment of these GO categories in the female CD/AP mice. To determine whether rhythmic proteins exhibit coordinated timing across functional pathways from any of the data sets, we next applied Phase Set Enrichment Analysis (PSEA) using the MSigDB M5 Gene Ontology Biological Process database to circadian proteins within each of the conditions (**Figures 5** and **S5**).^69–71^ PSEA identifies groups of biologically related proteins that are temporally coordinated over the circadian cycle. Across the male (LM, CD, AP, and CD/AP) and most of the female (LM, CD, and AP) conditions, we observed little coherent enrichment, with limited enrichment beyond cell-cycle-related terms (**Figures 5A, 5B** and **S5**). When compared to the noted PSEA enrichment in young males, this suggests, and agrees with previous investigations, that aging has a negative effect on GO pathway coordination in macrophages. ^19,72^

**Figure 5.**
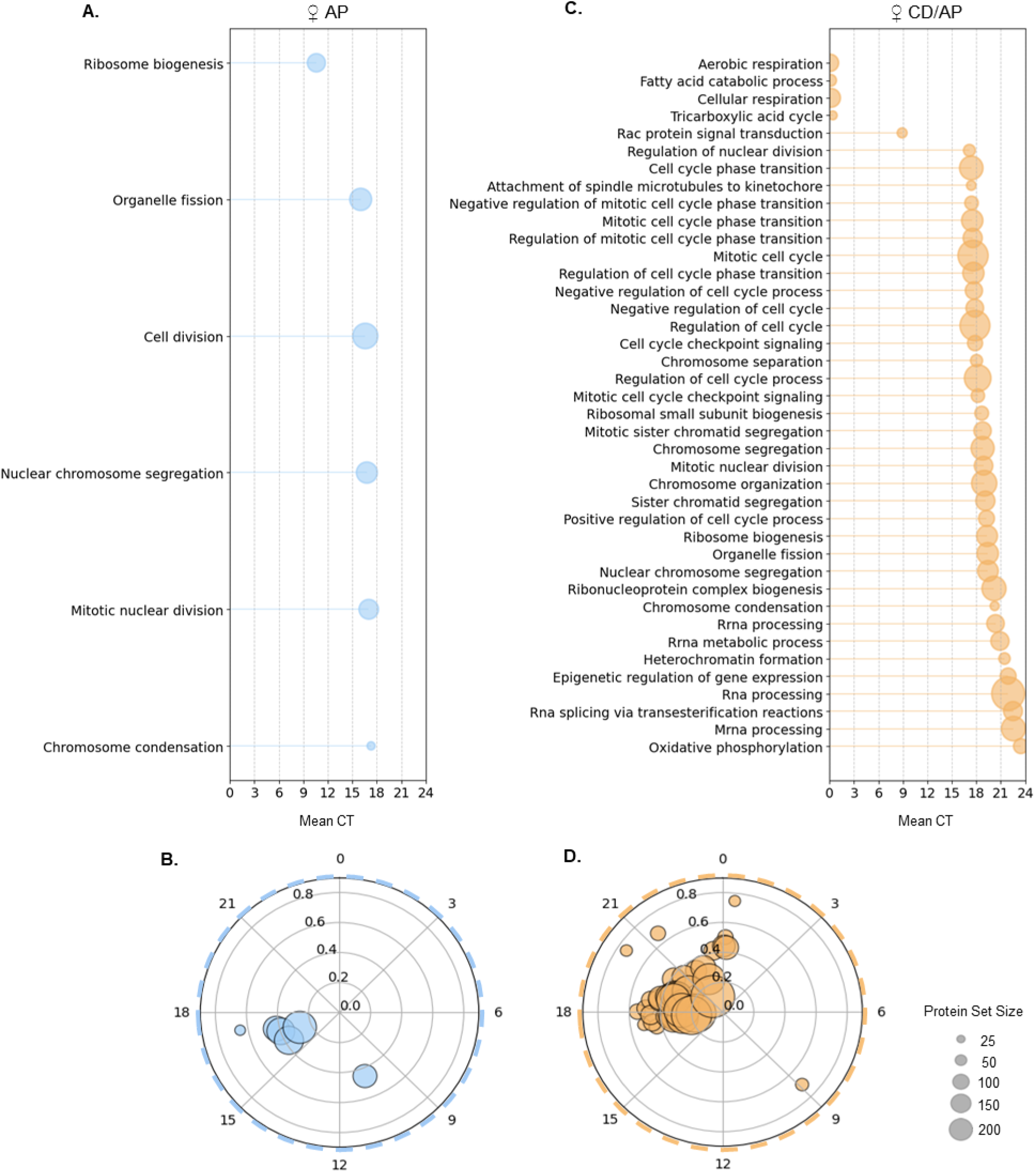
CD phase aligns gene ontologies involved in immunometabolism in the AP Female Macrophages. Circular scatterplot of significantly enriched GO pathways (Kuiper q ≤ 0.05) from Phase Set Enrichment Analysis (PSEA) of all rhythmic proteins (A) AP and (B) CD/AP female (♀) macrophages. Each point represents an enriched biological process, plotted by its vector-average circadian phase (angle, 0–24 h) and coherence magnitude (radius). The color denotes the macrophage group, and the point size scales with the number of genes within each pathway. (C) AP and (D) CD/AP linear “lollipop” plots showing mean circadian phase (0–24 h) for all significant pathways. Lines extend from 0 h to the pathway’s mean CT. Circle size indicates the number of unique proteins assigned to each GO term/functional category. Pathway labels correspond to GO Biological Process terms. Radial outlines denote sex (dashed = ♀) and color indicates lighting comparison (light blue = AP, orange = CD/AP). See also Figure S5.

In female CD/AP macrophages, PSEA analysis of all rhythmic proteins revealed coherent temporal enrichment of 30 GO Biological Process pathways (**Figure 5C**). Of these, we observed cell-cycle related terms but also other significantly coordinated pathways **(Figure 5C).** In particular, we saw additional clock coordination of immunometabolic pathways such as the TCA cycle and oxidative phosphorylation (aerobic respiration) that were tightly coordinated to peak in the late circadian night toward relative dawn **(Figure 5D)**. These findings indicate that female CD/AP macrophages have a tight phase-coordination of functional pathways for processes that can heavily influence the immune response, specifically immunometabolism and inflammation. The synchronized GOs here mirror the GOs of the phase-advanced proteins from the CD/AP females (**Figure 4**), suggesting that CD induces a phase-advancement of specific proteins that contributes to an overall alignment of pathways involved in the immunometabolic response, which may lead to enhanced inflammatory states.

### CD Shifts the Timing of Peripheral Macrophage Membrane Potential and Reactive Oxygen Species Generation

Given the significant alterations in the BMDM oscillatory proteome in CD/AP females, we next investigated how these changes affected macrophage functional outputs. To do so, we evaluated Aβ42 phagocytosis, mitochondrial membrane potential (ΔΨm), and intracellular reactive oxygen species (ROS) in synchronized BMDMs every 6 hours over 24 hours, in triplicate, using previously established methods (**Figures 6A-C** and **Table S7**).^18,19,65^ Here we identified something as rhythmic if it had an ECHO modeled period of 18-30 hours, and included overexpressed and repressed proteins, as the data was noisier and the sampling resolution lower. We found rhythmic Aβ42 uptake in male LM macrophages, in concordance with data previously gathered from young males.^18^ However, all other groups were determined to be arrhythmic by ECHO (**Figure 6A**). While interesting, the effect on phagocytosis did not align with the sex-specific effects of CD and AD, suggesting the loss of rhythmic phagocytic activity cannot explain the incongruity of reduced Aβ plaques in CD/AP females and the known negative effects of CD on AD in females.

**Figure 6.**
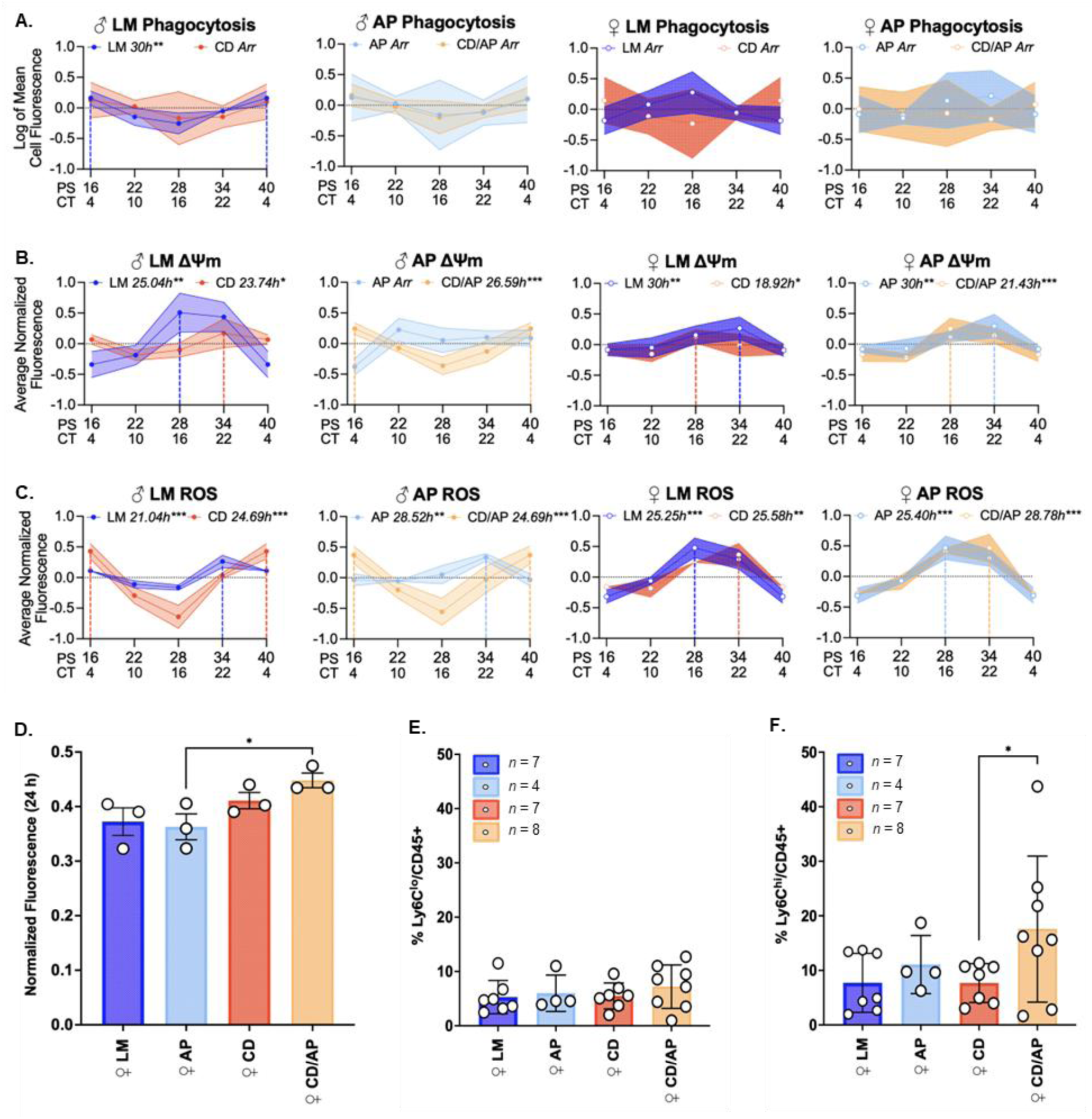
CD induces sex-specific changes that increase the inflammatory state of peripheral monocytes. (A–C) BMDMs from male (♂) and female (♀) LM, AP, CD, and CD/AP mice were synchronized in vitro by serum shock and sampled every 6 h over 24 h in triplicate. Plots show log-transformed fluorescence values representing Aβ42 phagocytosis (A), mitochondrial membrane potential (ΔΨm) (B), and reactive oxygen species (ROS) (C). Solid lines denote ECHO-modeled rhythmic fits, with shaded regions indicating ± 1SD of the normalized data at each time point. Data are displayed in post-shock (PS) time with inferred circadian time (CT).^16,19^ Period estimates (hours) are indicated where circadian rhythms were detected. Asterisks indicate significant ECHO fits (* p < 0.05, ** p < 0.01, *** p < 0.001, BH-adjusted). (D) Total mean ROS levels averaged across the 24-h sampling window (PS 16–40) in female macrophages (n = 3). Bars show mean ± SD with individual data points overlaid. (E) Circulating relative percentages of non-inflammatory (CD45⁺ Ly6Clo CD11b⁺) and (F) inflammatory monocytes (CD45⁺ Ly6Chi CD11b⁺) in female mice following 22 weeks of lighting exposure, identified using flow cytometry. Bars show mean ± SD with individual data points overlaid. Asterisks indicate significant group comparisons using uncorrected Fisher’s LSD (D–E) (* p < 0.05). Color indicates genotype/lighting condition (dark blue = LM, light blue = AP, red = CD, orange = CD/AP). ♂ = male, ♀ = female. See also Figure S6 and Tables S7-S9.

Contrary to phagocytosis, ΔΨm displayed robust rhythmicity across most conditions, with sex- and genotype-specific differences in period length and peak timing (**Figure 6B**). In LM male macrophages, ΔΨm oscillated with an approximately 24-hour period, with CD causing a delay in peak phase. While AP males were arrythmic, CD/AP males regained rhythmicity, though at a different peak timing to those rhythms seen in the LM or LM/CD. For females, ΔΨm circadianly oscillated, but CD consistently caused an advance in the peak phase of ΔΨm in both the LM and AP genotypes. Similar to ΔΨm, all ROS levels circadianly oscillated (**Figure 6C**). CD again altered the peak phase of ROS oscillations in a sex-specific manner. In males, CD produced a consistent delay in the peak phase of ROS. This delay of the peak phase of ROS was preserved in females, contrary to the advanced phases noted in ΔΨm for females. These phase relationships suggest parallel delays in the peak phases of ΔΨm and ROS in males due to CD, but a reversal of the peak phase timings of ΔΨm and ROS in females upon CD. This change in the phase relationship between ΔΨm and ROS in the CD/AD females suggests a decoupling of mitochondrial energetic state and redox output. Our results reveal the potential for sex-specific alterations in mitochondrial-redox coupling in AP BMDMs under chronic circadian disruption, which is consistent with a disrupted immunometabolic response and heightened inflammation.^73,74^

### The Combination of CD and AD Increases the Pro-Inflammatory Signature of Blood Monocytes in Female Mice

Permissibility of the brain to non-central nervous system privileged cell populations is generally low, except under conditions of heightened inflammation, as observed in AD cases.^75^ Given the coordinated increased infiltration of macrophages into the brain with the sex-specific alterations in macrophage functions and rhythmic coordination of immunometabolic processes, which could push CD/AD females toward enhanced inflammatory states, we next asked whether the BMDMs from the female CD/AP condition exhibited an elevated inflammatory state. To do so, we quantified one of the markers for inflammation, ROS.^76^ To track the overall daily ROS levels, we averaged normalized relative ROS fluorescence across the 24-hour sampling window (PS16–40) (**Figures 6D**, **S6A**, and **Table S8**). A three-way ANOVA demonstrated no effects due to sex, genotype, or lighting alone, but did show significant interactions across multiple conditions, indicating that ROS regulation is fundamentally distinct between males and females across conditions (**Table S8)**.

Given the significant difference between sexes, we next performed separate two-way ANOVAs for each sex. In females, we found that lighting only exerted a significant effect on ROS levels in the AP background, with maximal levels observed in the CD/AP condition (**Figure 6D**). In contrast, male BMDM ROS levels were more complex, with a significant decrease in ROS levels in the CD/AP condition and peak ROS levels in the CD and AP alone conditions **(Figure S6A)**. Together, these findings reveal opposing effects of sex on the oxidative responses in male and female BMDMs, with an increase in inflammatory phenotypes in the female CD/AD BMDMs as characterized by increased ROS levels.

Given this female-specific CD/AP increase in ROS, we hypothesized that circulating immune cell populations in blood collected from female mice would also show an increase in inflammatory phenotypes. To test this, we quantified circulating immune cell populations in blood collected from female mice at the end of the lighting protocol (**Figures 6E-F** and **Table S9**). Whole blood was labeled with CD45 to identify leukocytes, CD11b for monocytes, and Ly6C to distinguish non-inflammatory (CD45^+^ CD11b^+^, Ly6C^lo^) from inflammatory (CD45^+^, CD11b^+^, Ly6C^hi^) monocyte populations (**Figures S6B-D**).^77^ We found that inflammatory monocytes (**Figure 6F**) were selectively expanded in the CD/AP condition as compared to non-inflammatory monocytes (**Figure 6E**), indicating that the CD/AP condition promotes peripheral inflammatory monocyte expansion in females. This data parallels the elevated ROS levels and the increased recruitment of peripheral macrophages to the brain in CD/AP females, supporting a coordinated peripheral-to-central inflammatory response for females under chronic CD in AD.

## Discussion

CD is increasingly recognized as a biological stressor with direct consequences on human disease, including AD, though the mechanisms behind this effect have not been thoroughly elucidated.^7,10,11^ In this study, we evaluated how CD interacts with sex and immune function in an AD-susceptible context. Using ∼8-month-old APP/PS1 mice, we found a sex-specific divergence in CD effects on Aβ plaque burden, with female mice exhibiting an increase in baseline Aβ plaque pathology over males, consistent with the well-documented increase in AD incidence in females in both humans and mice.^27–29^ However, in females, CD reduced Aβ plaque pathology (**Figure 1**), a surprising outcome given that both human circadian disruption (e.g., shift work, irregular sleep-wake rhythms)^8,10,11,78^ and rodent CD paradigms^4,6,7^ generally worsen AD-relevant outcomes, including inflammation, cognitive decline, and neurodegenerative progression.

To investigate this paradox, we examined the sources of Aβ plaque clearance. Because microglia are the primary resident myeloid cells that surveil and respond to Aβ, and show clear sex difference, we were surprised to find microglial numbers showed only modest differences (**Figure 2**).^25–27,79^ Instead, we observed an increase in infiltrating macrophages that corresponded with the reduced Aβ plaque burden in CD/AP females. This aligns with evidence that, despite representing only ∼6% of plaque-associated myeloid cells, monocyte-derived macrophages disproportionately influence plaque clearance and plaque-proximal inflammation.^62^ Our results therefore add to the evidence of a role for peripheral macrophages in AD-relevant pathology.

While the relationship between macrophage infiltration and reduced Aβ burden supported the possibility that peripheral immune cells link CD to plaque modulation, these brain-resident and infiltrating populations did not fully resolve the paradox of decreased Aβ plaque burden under an environmental condition known to exacerbate AD pathology. To address this, we examined peripheral macrophage physiology from AP and CD/AP males and females, alongside littermate controls. In males, oscillations in macrophage behavior and clock-regulated proteins were largely maintained, with only a modest shift in protein peak timing and a decrease in ROS levels in CD/AP males compared to AP males (**Figures 3, 4** and **5**), suggesting minimal disruption to male immune-metabolic rhythms (**Figure 7**). Meanwhile, female peripheral macrophages from the CD/AP condition displayed a marked increase in the phase coordination of oscillatory proteins in metabolic and immunometabolic pathways (**Figure 3**), indicating a CD-induced reorganization and coordination of inflammatory timing specific to females (**Figure 7**). Mitochondrial-redox rhythms paralleled this sex- and CD-dependent reorganization, as while the temporal relationships of ΔΨm and ROS was largely preserved in males, these temporal relationships diverged in females specifically in the CD/AP condition (**Figure 6**). This loss of canonical ΔΨm-before-ROS coupling suggests a loss in antioxidant buffering and altered bioenergetic-redox coordination.^73,74^ This coupling aligned with both elevated daily ROS production and an expansion of inflammatory monocytes in the female CD/AP population (**Figure 6**).

**Figure 7.**
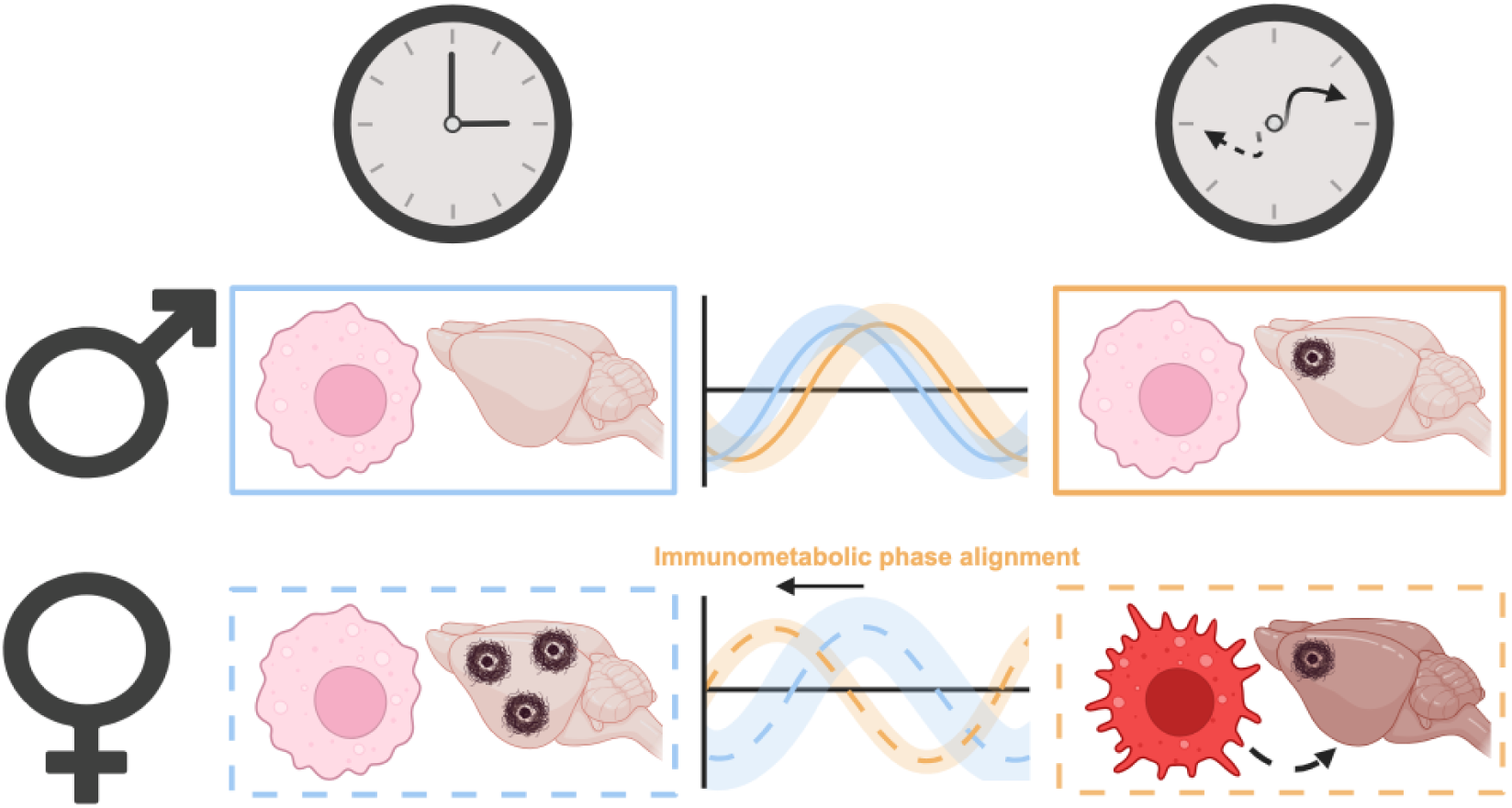
Chronic circadian disruption reprograms peripheral macrophage rhythms and alters early AD in a sex-specific manner. A diagrammatic summary of the effects of CD on macrophage circadian function and early AD-related pathology. Standard lighting conditions are depicted by a normal clock (top left), while the CD condition is represented by a distorted clock with misaligned hands (top right). Pictorial representations of male (♂; top row; solid lines) and female (♀; bottom row; dashed lines) APP/PS1 (AP) mouse macrophages, brains, and circadian proteomic oscillations are shown under standard and disrupted lighting conditions (CD/AP). Macrophages and brains appear pink when naïve and red/darker with increased inflammation states. The black dashed arrow indicates peripheral macrophage trafficking toward the brain. Brown tangles represent relative amyloid-β plaque burden. Representative circadian oscillations illustrate average phase relationships and CD-induced phase shifts. Arrows above the oscillations denote the direction and relative magnitude of phase change under CD/AP conditions compared with AP alone. The width of the shaded traces represents the alignment of the phases of the proteins within immunometabolic pathways. “Immunometabolic phase alignment” indicates increased temporal alignment of immunometabolic pathways in female CD/AP macrophages.

Regarding the paradox of lower Aβ plaque pathology and worsened disease phenotypes, our data suggest that CD reorganizes mitochondrial and immune-metabolic programs in female macrophages with AD pathology toward a more pro-inflammatory state, a pattern that has been shown in other contexts to potentiate tissue damage and neurodegenerative vulnerability.^7,26,80^ We hypothesize that CD reduces Aβ plaque burden in AP females via the increased infiltration of peripheral macrophages due to inflammatory signals. However, these peripheral macrophages have a heightened inflammatory state (reflected by increased ROS levels and cell surface markers) as compared to the non-CD population. Increased inflammatory activation is strongly implicated in accelerating neurodegeneration, cognitive decline, and tau pathology in both humans and mice (**Figure 7**).^6–9,25^ Thus, a plausible model is that CD transiently promotes Aβ clearance while simultaneously heightening inflammatory stress, ultimately increasing neuronal vulnerability.

Combined with the ability of inflammatory modulators to move back and forth across the BBB to control neuroimmune dynamics, our data supports evidence that the peripheral immune system plays a role in the development and progression of AD. Our data also aligns with the inconsistencies of the role for Aβ in AD, suggesting the importance of inflammation in the development of AD endophenotypes.^81,82^ Moreover, it suggests that CD affects the known circadian timing of the peripheral immune system to shape the sex-specific effects of Aβ pathology.^6,9,21,22,80^ Though systemic circadian stressors, including shift work and social jetlag, have emerged as epidemiological risk factors for dementia, and disrupted neuroimmune regulation are known to have sex-specific effects, the mechanistic links between the risks and the disease are poorly understood. ^5,7,10,11,13^ By revealing the sex-specific sensitivity of peripheral immune-metabolic rhythms to CD, our study identifies circadian–immune interactions as a potentially tractable method for mitigating AD risk, as well as a possible target for therapeutic treatment ^83,84^.

## Limitations of the study

Several limitations of this study warrant consideration. Although we observed reduced early amyloid plaque burden in CD/AP females, longitudinal studies will be required to determine if this reduction is sustained. We also did not directly trace the origin of infiltrating macrophages. Increased mortality in the APP/PS1 females necessitated rotation of wheel-running animals, which may have introduced variability in our behavioral analysis. Tissue limitations prevented co-staining of the same sections for amyloid-β, microglia, and macrophages. Finally, experimental throughput was constrained by the limited number of green-light circadian chambers, requiring staggered experimental rounds.

## Supporting information

All Supplemental Figures and Tables

## Resource availability

### Lead contact

- Requests for further information and resources should be directed to and will be fulfilled by the lead contact, Jennifer Hurley.

### Materials availability

- This study did not generate new unique reagents.

### Data and Code Availability

- Proteomics data will be deposited at ProteomeXchange and be publicly available as of the date of publication.
- All supplementary data have been deposited at Mendeley Data (preview at https://data.mendeley.com/preview/kh5g2zwcmm?a=907c08fe-e43e-4b57-ae27-969891529a58) and will be publicly available as of the date of publication.
- This paper does not report original code.
- Any additional information required to reanalyze the data reported in this paper is available from the lead contact upon request.

## Acknowledgements

We thank Mariana Figueiro, Sagan Legett, Lily Donaldson, Robert Karlicek, and Barbara Plitnick for guidance during the lighting setup and for sharing samples. We thank Dani Gullick, the Neural Stem Cell Institute, the RPI Microscopy Core Facility, and Cameron Plowinske for help with imaging. We acknowledge the RPI Bioresearch Core facilities, for animal care and technical support. This work was supported by a grant from the NIH-NIGMS to J.M.H. (R35GM128687) and a gift from the Warren Alpert Foundation to J.M.H. and J.S.D.. C.V.C., N.G.F., G.T.C., and J.R.C. were supported by a NIH-NIA T32 training grant (T32AG078123 and T32AG057464). N.G.A., E.J.C. and M.S.J were supported by an NIH-NIGMS T32 training grant (T32GM141865 and T32GM067545).

## Author Contributions

J.M.H., J.S.D., M.S.J., C.V.C., and G.T.C. conceptualized the research. C.V.C., J.M.H., and M.S.J. curated the data. C.V.C., J.R.C., N.G.F., G.T.C., N.G.A., A.A.H., M.L., and E.J.C. – performed formal analysis. J.M.H. and J.S.D. acquired funding. C.V.C., N.G.F., G.T.C., N.G.A., and E.J.C. performed experiments. J.M.H., C.V.C., M.S.J., J.R.C., G.T.C., R.U., A.A.H., M.L., E.J.C., and M.B. developed methodology. J.M.H., J.S.D., C.V.C., and M.S.J. were responsible for research project administration. J.M.H., J.S.D., and J.R.C. provided study material resources. J.R.C.-implemented software. J.M.H. and J.S.D. provided supervision. C.V.C., M.S.J., J.R.C., and N.G.F. validated the reproducibility. C.V.C., N.G.F., N.G.A., and J.M.H. prepared the visualization of the data. C.V.C., J.M.H., M.S.J., N.G.A., and G.T.C. prepared the original draft writing. J.M.H., N.G.F., and M.S.J. edited the paper.

## Materials and Methods

### Key resources table

| REAGENT or RESOURCE | SOURCE | IDENTIFIER |
| --- | --- | --- |
| Antibodies |  |  |
| Anti-Aβ 6E10 (rabbit monoclonal) | Invitrogen | #MA5-48043 |
| Anti-IBA1 (rat monoclonal) | Invitrogen | #MA5-38266 |
| Anti-TMEM119 (rabbit polyclonal) | Invitrogen | #PA5-119617 |
| Donkey anti-rat IgG Alexa Fluor 647 | Invitrogen | #A48272 |
| Donkey anti-rabbit IgG Alexa Fluor 568 | Invitrogen | #A10042 |
| ProLong Gold Antifade Mountant with DNA stain DAPI | Invitrogen | #P36931 |
| Anti-CD45 (clone 30-F11) | Invitrogen | #14-0451-82 |
| Anti-CD11b (clone M1/70) | Invitrogen | #14-0112-82 |
| Anti-Ly6C (clone HK1.4) | Invitrogen | #45-5932-82 |
| Critical commercial assays |  |  |
| Luminex Milliplex Mouse Cytokine/Chemokine 32-plex magnetic bead panel | Millipore | MCYTMAG-70K-PX32 |
| JC-10 Mitochondrial Membrane Potential Assay | Abcam | #ab112134 |
| DCFDA ROS Detection Assay | Abcam | #ab113851 |
| CyQUANT Cell Proliferation Assay | Invitrogen | Cat# C7026 |
| HiLyte Fluor 488-labeled Aβ42 | AnaSpec | #AS-60479-01 |
| Deposited Data |  |  |
| Supplementary Data | Mendeley Data | See Data Availability Statement |
| Proteomics Data | ProteomeXchange | See Data Availability Statement |
| Experimental models: Organisms/strains |  |  |
| APPswe/PS1dE9 (APP/PS1) mice | Jackson Laboratory | Stock #034829 |
| C57BL/6J mice | Jackson Laboratory | Stock #000664 |
| Software and algorithms |  |  |
| STARR Life Science In cage running wheels | Starr Life Sciences | Model VV-ICRW-W/SNSR |
| Actimetrics Wireless Low-profile Running Wheels | Actimetrics | Model ACT-557-WLP |
| VitalView software | Philips/Respironics | Version 5.1 |
| Actimetrics ClockLab Data Collection System | Actimetrics | Model AM1-CD01 |
| Actimetrics ClockLab Analysis Package | Actimetrics | Model AM1-CA06 |
| TissueFAXS Q | TissueGnostics | Version 8.0.76 |
| StrataQuest | TissueGnostics | Version 8.0.76 |
| ECHO | Github/delosh653/ECHO | Version 4.0 |
| LIMBR | Github/aleccrowell/LIMBR | Version 0.2.10 |
| PSEA | Github/ranafi/PSEA | Version 1.1 |
| Prism | GraphPad | Version 10.1.0 |
| MATLAB | MathWorks | Version R2023a |
| Python | Python Software Foundation | Version 3.12.12 |
| R | R Core Team | Version 4.4.2 |
| RStudio | Posit Software, PBC | Version 2024.09.1+394 |
| R package: org.Mm.eg.db | Carlson, M. (2019) | Version 3.2<br>Bioconductor |
| MSigDB M5 (GO Biological Processes) | Broad Institute | Mouse, M5 |
| Other |  |  |
| 525 nm green LED lighting (4 $\mu\text{W}/\text{cm}^2$) model and scheduler | Custom | Scheduler LE365S-41 |
| Orbitrap Eclipse Tribrid Mass Spectrometer | Thermo Fisher Center for Multiplexed Proteomics at Harvard Medical School | Eclipse Tribrid MS (Catalog # FSN04-10000), coupled with EASY-nLC 1200 system |
| LSRII flow cytometer | Becton Dickinson | TA Instruments TGA-Q50 (Rensselaer CBIS Analytical Biochemistry Core Facility) |

### Ethics statement

All experiments were performed in accordance with the National Institutes of Health Office of Intramural Research and were approved and supervised by the Rensselaer Polytechnic Institute Animal Care and Use Committee (protocol numbers HUR-001-21, HUR-003-21, and HUR-001-24).

### Animal models and lighting protocols

#### Simulated standard and disruptive lighting for APP/PS1 Alzheimer’s disease model and littermate controls

Hemizygous APPswe/PS1dE9 (APP/PS1) mice and non-transgenic littermate controls were obtained from Jackson Laboratory. Male and female mice (8–10 weeks old) were acclimated under standard 12:12 h light–dark (white light) conditions for 2 weeks before transitioning to monochromatic green light (525 nm LEDs, 4 μW/cm²).^44^ Mice were assigned to either standard lighting (12 h light : 12 h dark) or circadian disruption (CD) consisting 4 days 12L:12D followed by 3 days 12D:12L, repeated for 22 weeks. Mice were group-housed unless single housing was required for behavioral assays.

#### Estrus cycle synchronization

To induce the Whitten effect, male soil bedding was distributed in the cages of female mice four days prior to collection time points to expose them to male pheromones and induce the estrus cycle.^40^ This approach synchronized their estrus cycles and allowed for sample collection after peak estrogen levels, thereby minimizing hormone variability.^39,40,85^

### Behavioral analyses

#### Wheel running activity

Subsets of mice were individually housed with running wheels throughout the 22-week protocol. Wheel rotations were continuously recorded at 10-min intervals using VitalView (half of all males) or ClockLab (half of all males and all females). Circadian period and interdaily stability were calculated using chi-squared periodograms. Outliers were identified using a Dixon Q test (p < 0.05) and excluded. Statistical analysis of activity levels over time can be found here.^86^

#### Nesting behavior

At baseline, week 11, and week 22, mice were individually housed with four pre-weighed cotton nestlets. After 24 h, the nests of the first fourteen mice in each group were photographed and weighed. Nest quality was scored by two blinded observers using established criteria, and nestlet usage was calculated as the change in nestlet mass.^49^

### Brain tissue processing and immunohistochemistry

#### Sectioning and immunostaining

Brains were fixed in 4% PFA (Thermo Fisher #28908), cryoprotected in 15% and 30% sucrose, embedded in OCT (Thermo Fisher #23730571), and sagittally sectioned from the center at 25 µm. Free-floating sections were permeabilized (0.5% Triton X-100), blocked (5% normal donkey serum (Abcam #AB7475), 3% bovine serum albumin (Thermo Fisher #BP1600-100), 0.3% Triton X-100) for 2 h. Sections were incubated overnight at 4°C with primary antibodies (6E10, IBA1, TMEM119), and subsequently with AlexaFluor secondary antibodies and DAPI after mounting.

#### Imaging and Quantification

Images were acquired using a TissueFAXS Q confocal system (20×/0.8 NA objective, 16-bit sCMOS camera), with each field focused at the plane of highest signal. Quantification of plaques and microglia/macrophages was performed using StrataQuest (v8.0.76) using fixed thresholds per round. Aβ-positive areas were identified by selecting the brightest spots within a pixel intensity range of 3500–12,000 (Cy3 channel). Plaque area was normalized to the region of interest. Microglia and macrophages were detected using nuclear masks generated from DAPI images and surrounding cytoplasmic signal (6–10 µm expansion) was assessed for TMEM119 (microglia-specific) or IBA1 (microglia/macrophages). Empirical intensity cutoffs were applied to classify cells. Outputs included cell density and plaque-associated microglia percentages. To validate antibody specificity, bone marrow-derived macrophages (BMDMs) were fixed, stained with IBA1 and TMEM119, then imaged on a Nikon Ti2 microscope equipped with a CFI Apo 60× 1.49 NA objective using identical exposure and gain settings. Sex × genotype × lighting interactions were performed using a three-way ANOVA. Significant interactions were further examined by two-way ANOVA followed by Tukey’s post-hoc tests. Statistical significance was defined as *p* values are indicated as * <0.05, ** <0.01, *** <0.001, **** <0.0001. Statistical analyses were performed in GraphPad Prism. Outliers were identified using the interquartile range (IQR) method.

### Blood and immune cell analyses

#### Serum cytokine profiling

Serum was collected from tail vein bleeds at weeks 1, 11, and 22, centrifuged at 10,000 × g for 10 min, and stored at −80 °C until analysis. Cytokine concentrations were quantified using a 32-plex Millipore Luminex magnetic bead panel (MilliporeSigma #MCYTMAG-70K-PX32) according to the manufacturer’s instructions. Samples were incubated with capture beads, detection antibodies, and streptavidin-phycoerythrin. Standard curves and quality control beads were included on each plate to ensure assay performance. Mean fluorescence intensities were converted to pg/mL and averaged across technical duplicates. Longitudinal changes were analyzed using linear mixed-effects models, and between-group comparisons were performed using Welch’s t-tests with Sidak correction for multiple comparisons in GraphPad Prism.

#### Inflammatory monocyte analysis

At study endpoint, whole blood was collected via cardiac puncture. Leukocytes were Fc-blocked and stained with antibodies to CD45, CD11b, and Ly6C.^77^ Flow cytometry was performed on an LSRII cytometer. Prior to statistical analysis, mouse weights and liver weights were screened for outliers using the ROUT methods (Q = 5%). Two mice were removed due to weight outliers (Round 2, ear tags AP 29 and CD/AP 12), followed by removal of three mice as proportional monocyte measurement outliers (Round 2, ear tag CD/AP 13; Round 3, ear tags LM 55 and CD 41). Identified outliers were removed from further analysis. Within each experimental group (male and female LM, AP, CD, CD/AP), monocyte data were also screened for outliers using ROUT (Q=5%). Two-way ANOVA evaluated effects of lighting and genotype, with Fisher’s LSD for post-hoc comparisons in GraphPad Prism.

### Ex vivo macrophage assays

#### Macrophage isolation and synchronization

BMDMs were isolated from femurs and tibias of mice at endpoint, differentiated in Dulbecco’s Modified Eagle’s Medium (DMEM; 4.5 g/L glucose, L-glutamine, sodium pyruvate), and grown to confluency in 35 mm cell plates following established protocols.^18,19^ To synchronize circadian rhythms, cultures underwent 24 h serum starvation followed by a 2 h serum shock (50% fetal bovine serum) and were then returned to Leibovitz’s L-15 medium containing luciferin and 10% fetal bovine serum. Circadian assays were initiated ≥16 h post-shock to allow synchronized cells to re-establish homeostasis.

### Aβ42 phagocytosis assay

Serum-synchronized BMDMs were exposed to 0.25 mg/mL HiLyte Fluor 488–labeled Aβ42 as previously described every 6 h for 24 h.^18^ After each time point, cells were collected, fixed, then imaged on a Zeiss LSM 510 confocal microscope using a 40X objective. Pixel intensities from 110 randomly selected cells per time point were quantified in MATLAB. Intensities were log10-transformed, IQR outliers were removed, and rhythmicity was assessed using ECHO ^65^. This randomization was performed 10 times per condition and sex, and the averaged ECHO model parameters were reported.

#### Reactive oxygen species (ROS) and mitochondrial membrane potential (ΔΨm) analysis

Naïve BMDMs were plated in 96-well plates (8,000 cells/well) and allowed to adhere and grow to confluency. Cells were then serum-synchronized using the same serum synchronization protocol described above, scaled for 96-well format. For ROS, cells were incubated with DCFH-DA (20 µM) for 45 min and fluorescence was read at Ex/Em = 485/535 nm and normalized to CyQUANT cell counts on a Tecan i-control infinite 200 Pro version 2.0.10.0 plate reader. ΔΨm assays were performed using JC-10 assays per manufacturer instructions; fluorescence was measured at Ex/Em = 490/525 nm and 540/590 nm on a Tecan i-control infinite 200 Pro version 2.0.10.0 plate reader. Values were normalized by subtracting background readings from wells with unstained cells in assay media and from cell-free wells. The ratio of 590/525nm was calculated and technical replicates were averaged denoting biological replicate. Daily averages and rhythmicity were evaluated using ECHO. Statistical analyses used three-way ANOVA for sex × lighting × genotype, followed by sex-stratified two-way ANOVA when interactions were present, using GraphPad Prism.

### Proteomics

#### Sample preparation and LC–MS/MS analysis

BMDMs from male and female mice were collected every 3 h across a 24 h cycle, with three independent biological rounds per sex. Lysis, protein extraction, reduction, alkylation, tryptic digestion, and labeled using TMTpro 18-plex reagents were performed at the Harvard Center for Mass Spectroscopy. Peptides were fractionated and analyzed on a Thermo Orbitrap Eclipse with EASY-nLC 1200, 120-min gradient, full MS at 120,000 resolutions, using the Synchronous Precursor Selection (SPS) MS^3^ quantification. Peptide-spectrum matches (PSMs) were assigned using an in-house Sequest-based algorithm, and identifications were filtered to a 1% false discovery rate (FDR). TCMP provided peptide-level reporter ion data that was summed across repeated identifications, normalized across TMT channels, and log2-transformed to reduce skew while maintaining their proportional representation, and mapped to UniProt accessions.^87,88^

#### Data processing and rhythm detection

Batch correction and imputation of peptide-level signal-to-noise (S/N) values were performed using the LIMBR pipeline with default parameters.^64^ LIMBR was applied separately to each genotype × lighting treatment to avoid bias. Proteins with >30% missing values across timepoints were excluded. Protein-level abundances were calculated by averaging S/N values across all unique peptides per protein. Rhythmicity was detected using ECHO in free-run mode, applying smoothing, normalization (z-scoring), and linear detrending.^65^ Core clock proteins were visualized as mean ± standard deviation at each time point using GraphPad Prism. Circadian protein classification was defined for proteins detected in at least 70% of time points across all datasets with the following ECHO parameters: (a) Benjamini–Hochberg (BH) adjusted *p* < 0.05, (b) a period of 18-26 hours consistent with circadian oscillations determined from wheel running and core clock protein periods, and (c) rhythmic models corresponding to harmonic, forced, or damped oscillations. Peak expression times were converted to circadian time (CT, 0–24 h) and normalized using Hours Shifted × (24 / Period) + 12 to align with a 24 h rhythm. Twelve hours were added to align post-synchronization (PS) times with CT.^16,19^

#### Visualization of Circadian Data

Percent circadian proteins were visualized using GraphPad Prism. Temporal distributions of peak CT’s were visualized as radial histograms (circular density plots) in polar coordinates, with CT0 at the top proceeding clockwise, using Python. Heatmaps of relative normalized protein abundances were generated in RStudio using the ggplot2, patchwork, and cowplot packages after scaling values to be between -1 and +1 per for each protein. Scatter density plots and marginal histograms comparing CT distributions under standard versus disruptive lighting were produced in Python using pandas, numpy, matplotlib, and seaborn packages, with kernel density estimates calculated using a von Mises distribution (κ = 5). UpSet plots visualizing unique and shared circadian proteins across conditions were generated with the Python upsetplot package.

#### Functional Enrichment Analysis

Sex- and lighting-specific differences were assessed first by identifying proteins that were uniquely rhythmic in each condition (from Upset plots). Over-representation analysis (ORA) of uniquely rhythmic proteins was performed using clusterProfiler in RStudio, org.Mm.eg.db, and AnnotationDbi,^67,68,89^ with the full set of detected proteins in each dataset used as the background. All Gene Ontology (GO) categories for Biological Process (BP), Cellular Component (CC), and Molecular Function (MF) were assessed,^66^ but BP results were reported to highlight broader biological processes.

#### Phase shift and functional annotation

Circadian proteins were classified as phase-advanced (ΔCT < −3 h) or phase-delayed (ΔCT > +3 h). UniProt identifiers were mapped to MGI gene symbols using biomaRt in RStudio. Functional annotation employed GO Biological Process terms via clusterProfiler and UniProt keyword searches.^67^ Proteins were assigned to four curated functional groups: Immune (keywords: “immune”, “inflamm”, “cytokin”, “leukocyte”), Motility (“motil”, “migration”, “cytoskeleton”, “actin”), Mitochondrial (“mitochondr”, “respirat”, “ATP”, “electron transport”), and ROS/oxidative stress (“ROS”, “oxidative”, “peroxide”, “redox”). ΔCT contrasts were computed for each male and female disrupted versus standard lighting pair (CD – LM or CD/AP – AP), and GOBP enrichment was run separately for advanced, delayed, and in-phase categories using all circadian genes as the background. Functional group assignments informed visualization and interpretation in RStudio but were not used for statistical testing. Circular kernel density estimates were generated using a von Mises distribution (κ = 5). All violin plots were produced in Python 3.12.12 using pandas, matplotlib, seaborn, and scikit-posthocs.

#### Phase set enrichment analysis (PSEA)

Proteins exhibiting robust rhythmicity (ECHO period 18–26 h, BH-adjusted p < 0.05) were mapped from UniProt identifiers to gene symbols using AnnotationDbi and org.Mm.eg.db in RStudio.^68,89^ For each dataset, CT values were averaged across duplicate gene symbols to produce a single CT per gene for Phase Set Enrichment Analysis (PSEA) analysis, and the resulting gene–CT tables were saved as tab-delimited files for PSEA. PSEA analyses were then performed using the Java tool with the MSigDB M5 mouse GO Biological Process GMT file (m5.go.bp.v2025.1.Mm.symbols.gmt; circadian domain 0–24 h, minimum set size 10, 10,000 simulations, q < 0.05).^69^ Temporal clustering results were visualized in Python using pandas, numpy, matplotlib, seaborn, and upsetplot. Circular (polar) plots represented vector-average circadian time and magnitude. Lollipop plots depicted vector-average values per pathway with point size proportional to the number of genes.

