## Supplementary material for "Circadian disruption induces sex-specific Alzheimer’s pathophysiology and immune cell reprogramming": All Supplemental Figures and Tables

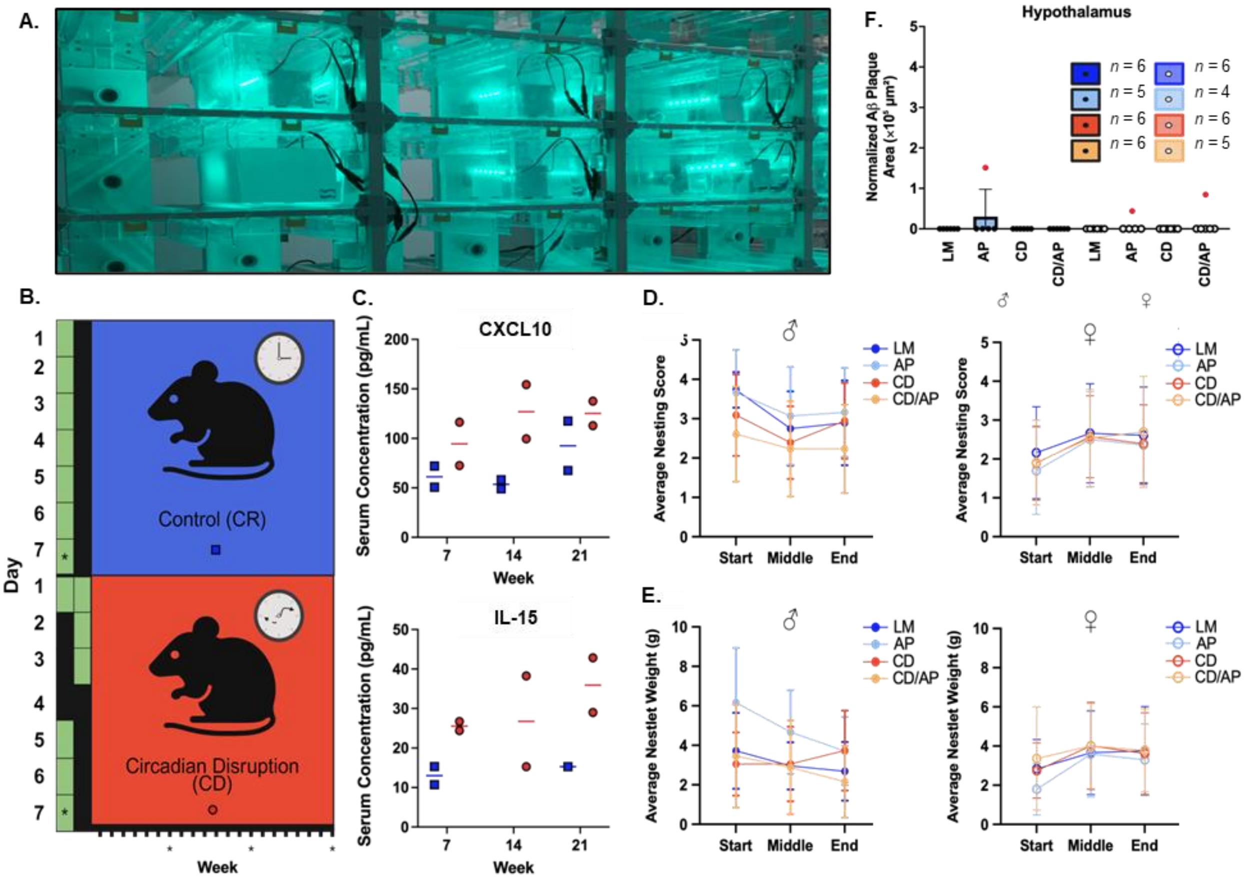


**Figure S1. CD protocol affects inflammatory serum cytokine levels but not behavior or Aβ accumulation in the hypothalamus. Related to Figure 1.**

(A) A representative image of the green lighting cages. (B) Experimental design of C57BL/6 male mice exposed to lighting schemes simulating either control (CR) or circadianly disruptive (CD) schedules used to gather preliminary data on serum cytokines (*n* = 2). Weekly lighting schemes are depicted with 12-hour increments of "lights on" (green) and "lights off" (black) for each day. Asterisks indicate the timing of blood draws for sampling, which were profiled every seven weeks over the 22-week protocol. (C) Serum cytokine levels of the pro-inflammatory cytokines CXCL10 and IL-15 reported in pg/mL. Each data point represents an individual mouse, with blue squares representing CR mice and red circles indicating CD mice. Horizontal lines denote the average serum concentrations. (D-E) Average nestlet scores (D) and weights (E) measured at the start, midpoint, and end of the lighting protocol in LM, AP, CD, and CD/AP male and female mice (*n* = 14). Data are presented as mean ± standard deviation of the mean. (F) Total Aβ plaque area in the hypothalamus normalized to the region of interest in male and female brains (*n* = 6 except in the case of the female AP hypothalamus where *n* = 5). Red dots represent outliers identified using the interquartile range (IQR) method. One male in the AP condition was identified as an outlier (38.898801 μm²) and was out of graphical bounds. This point is therefore not shown on the graph. Bars show mean ± SD. Dark blue with black outline = littermate (LM) male mice, light blue with black outline = APP/PS1 (AP) male mice, red with black outline = circadianly disrupted (CD) male mice, orange with black outline = CD/AP male mice, dark blue with blue outline = littermate (LM) female mice, light blue with blue outline = AP female mice, red with red outline = CD female mice, orange with orange outline = CD/AP female mice. ♂ = male mice, ♀ = female mice.

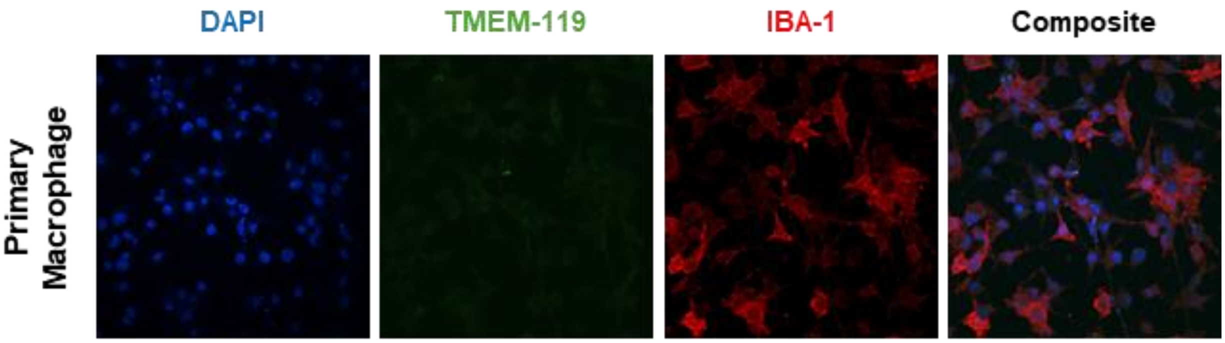


**Figure S2. Validation of antibody specificity of bone marrow derived macrophages *in vitro*. Related to Figure 2.**
Representative images of bone marrow derived macrophages (BMDMs) showing DAPI nuclear counterlabeling and TMEM119 and IBA1 immunoreactivity. Individual channels are shown separately, and a composite image displaying all three labels is also presented (white scale bar = 20 μm).

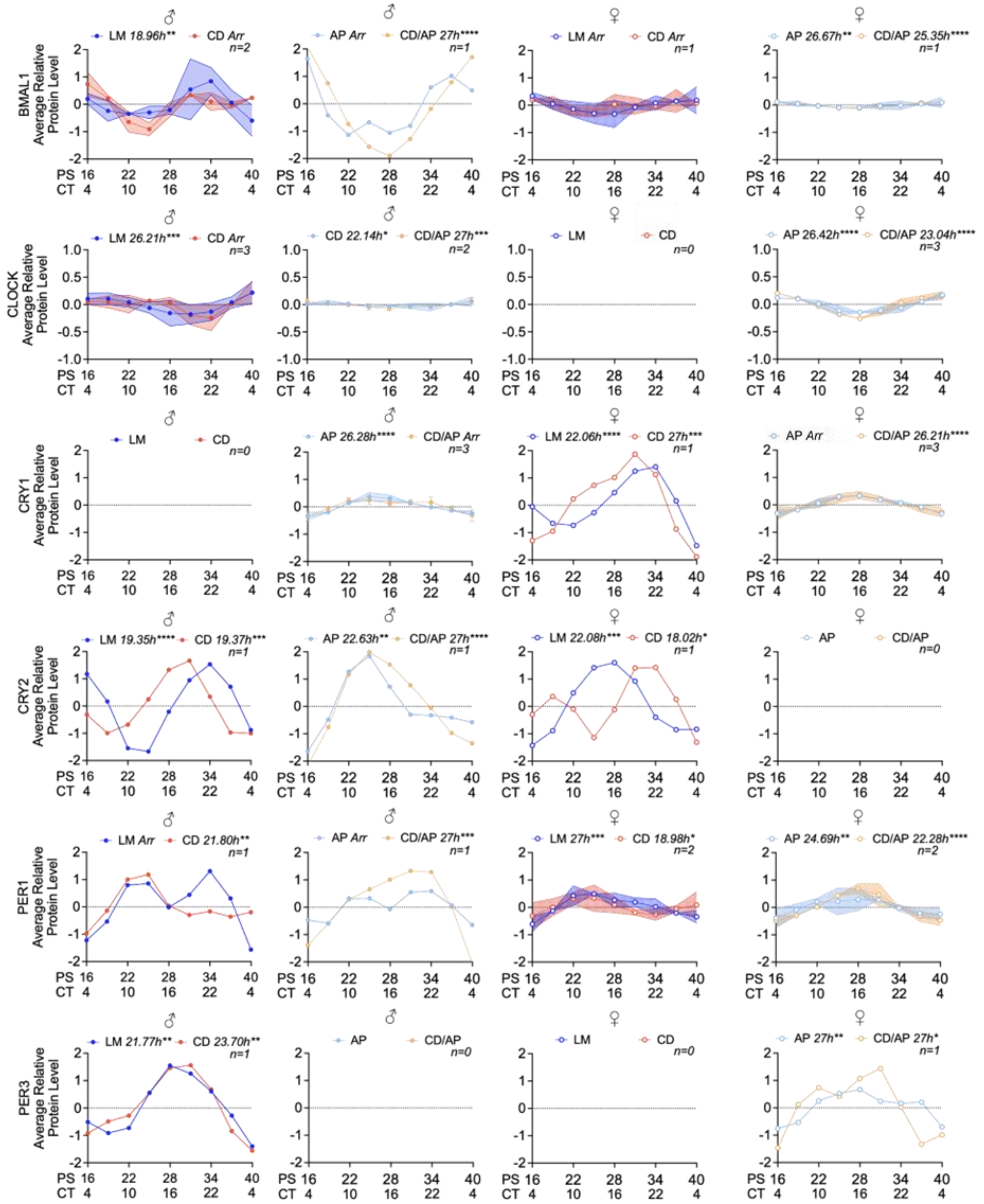


**Figure S3. Macrophage core clock proteins show sex-specific responses to CD and AP. Related to Figure 3.**

Average relative protein levels of the core clock proteins BMAL1, CLOCK, CRY1, CRY2, PER1, and PER3 from the male and female LM, AP, CD, and CD/AP treatment conditions every 6 hours over a 24-hour period. Plots lines show average of normalized, smoothed and detrended data for each protein, with shading representing ± 1 SD at each time point if the protein was detected in multiple time courses. The period of each rhythm (in hours) is indicated on the plots, with asterisks denoting statistically significant rhythmicity based on ECHO (* p < 0.05, ** p < 0.01, *** p < 0.001, BH-adjusted). Lines with closed points are male (♂) while open points are females (♀), and colors indicate genotype/condition: dark blue = LM, light blue = AP, red = CD, orange = CD/AP.

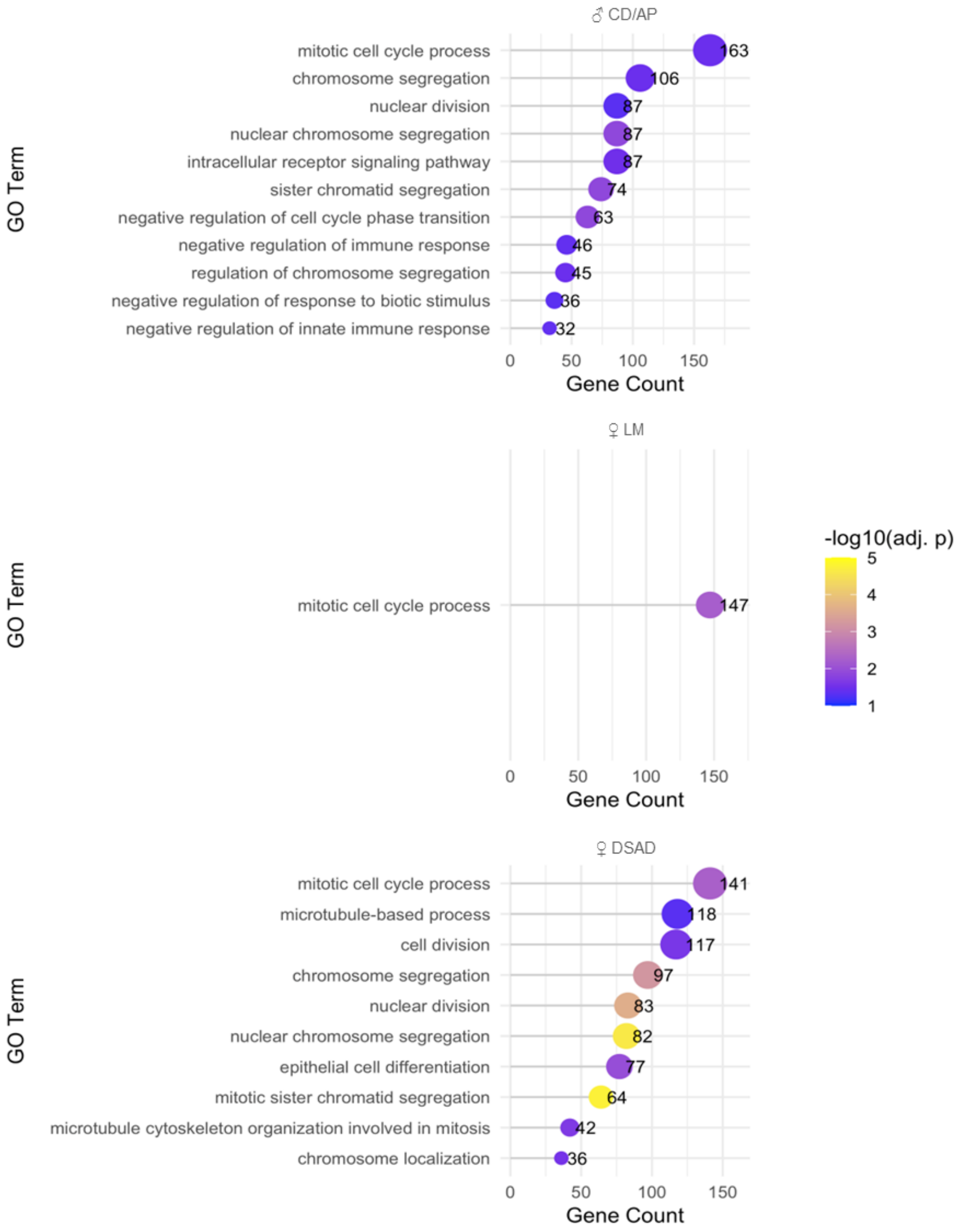


**Figure S4. Lollipop plots of enriched GO biological process terms in detected circadian proteins. Related to Figure 3.**

Top enriched Gene Ontology (GO) biological process (BP) terms for circadian proteins were identified using overrepresentation analysis (ORA). Only datasets with significant enrichment results are shown, specifically male (♂) CD/AP, female (♀) LM, and female

AP. For each term, the number of contributing genes is shown on the x-axis, and -log10(BH adjusted p-value) is represented by the color intensity of the dots. Grey lines connect each dot to the y-axis for clarity. GO terms were filtered for significance (FDR < 0.05) and simplified to reduce redundancy (semantic similarity cutoff = 0.7).

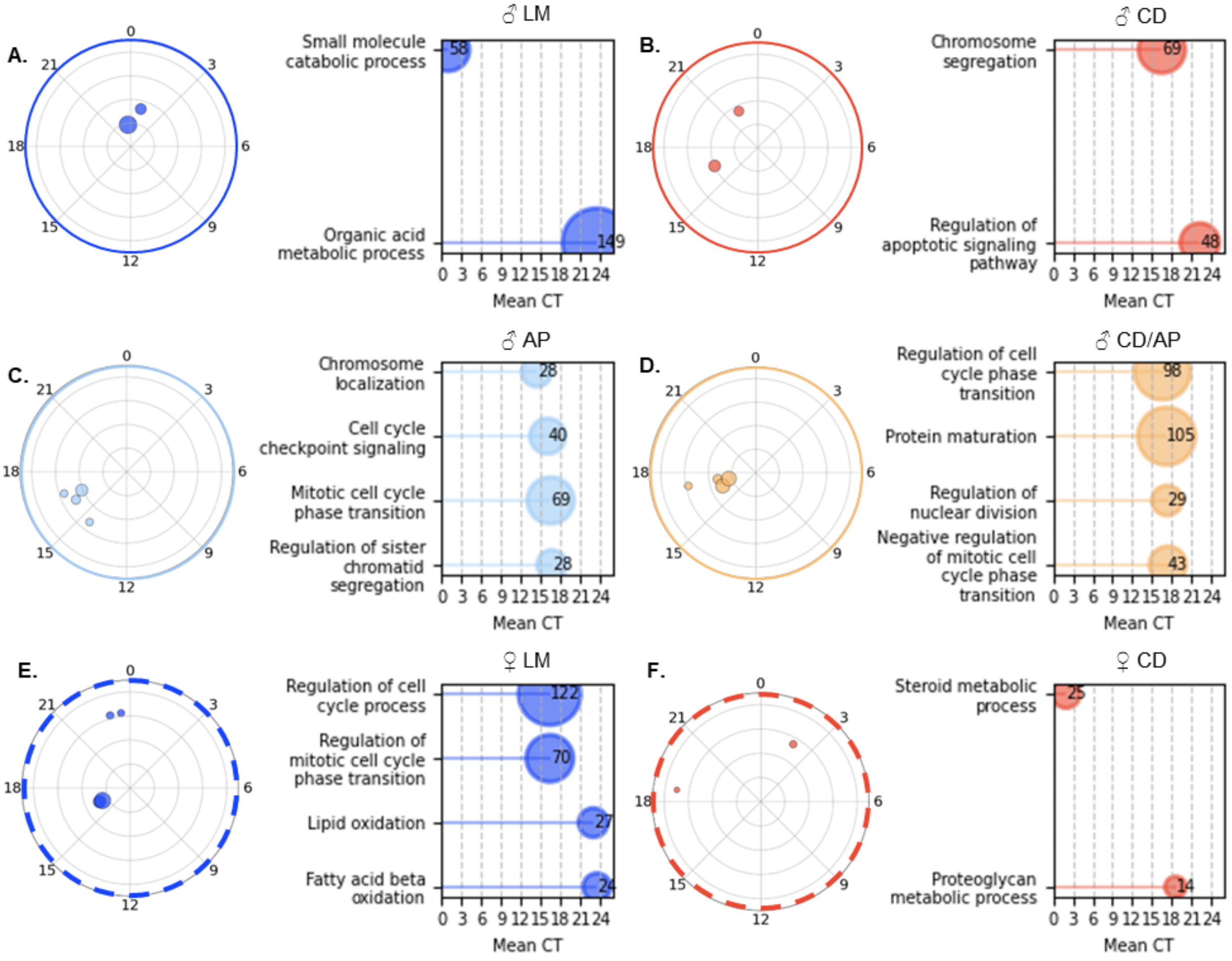


**Figure S5. CD imparts little phase alignment of gene ontologies in male and female macrophages beyond the AP background. Related to Figure 5.**

Circular scatterplot of significantly enriched GO pathways (Kuiper q ≤ 0.05) from Phase Set Enrichment Analysis (PSEA) of all rhythmic proteins in (A) LM male (♂), (B), CD male, (C) AP male, (D) AP/CD male, (E) LM female (♀), and (F) CD female macrophages. Each point represents an enriched biological process, plotted by its vector-average circadian phase (angle, 0–24 h) and coherence magnitude (radius). Color denotes the macrophage group, and the point size scales with the number of genes within each pathway. Linear lollipop plots showing mean circadian phase (0–24 h) for all significant pathways with lines extending from 0 h to the pathway's mean CT. Circle size indicates the number of unique proteins assigned to each GO term/functional category. Pathway labels correspond to GO Biological Process terms. Radial outlines (solid = ♂, dashed = ♀) and color indicates lighting comparison (dark blue = LM, light blue = AP, red = CD, orange = CD/AP).

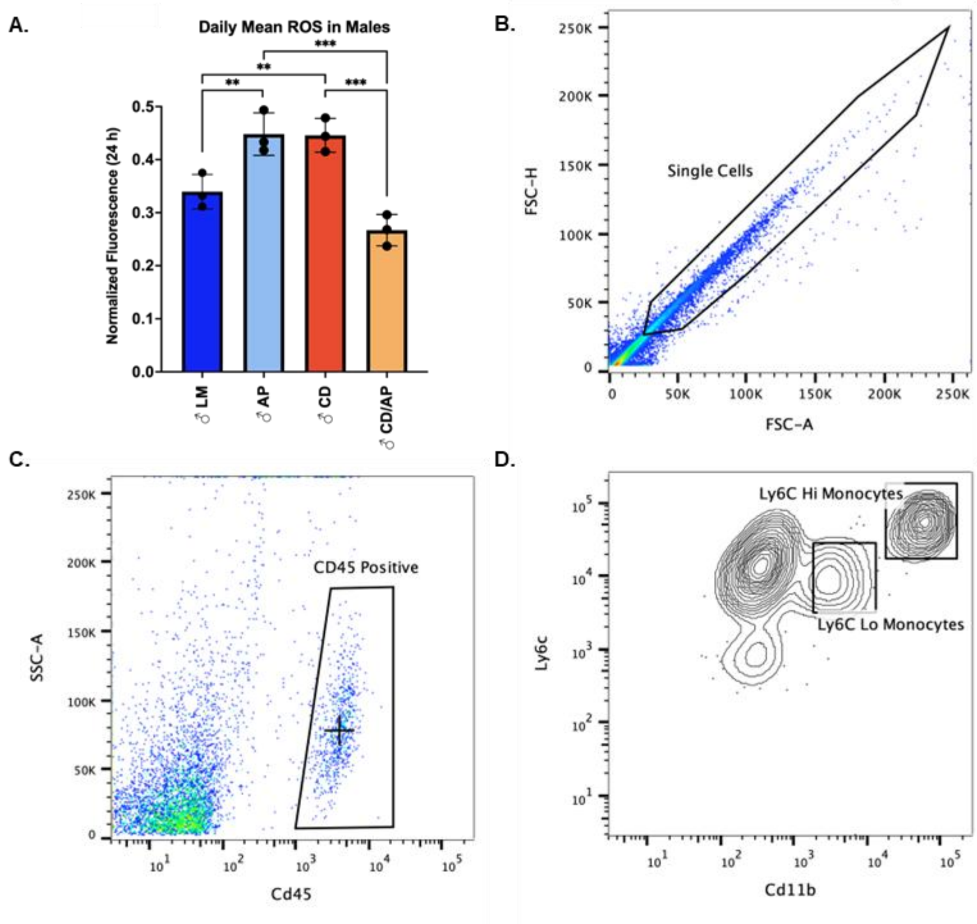


**Figure S6. Daily mean reactive oxygen species (ROS) is affected by CD. Related to Figure 6.**

(A) Daily mean reactive oxygen species (ROS) values averaged from all timepoints during the 24-hour sampling window (PS 16-40) for male (♀) mice (*n* = 3). Bars show mean ± SD with fill color indicating genotype/lighting condition (dark blue = LM, light blue = AP, red = CD, orange = CD/AP) with individual points overlaid. Multiple comparisons were performed using uncorrected Fisher's LSD; * $p$ = < 0.05. (B) Flow cytometry plot of forward-scatter height (FSC-H) vs forward-scatter area FSC-A used for doublet discrimination. Black box denotes flow cytometry gates used to identify singlets. (C) Flow cytometry plot of side-scatter area (SSC-A) vs cluster of differentiation 45 (CD45) identifying leukocyte populations. Black box denotes flow cytometry gates used to identify $CD45^{+}$ leukocytes. (D) Flow cytometry plot of lymphocyte antigen 6 complex, locus C (Ly6C) vs cluster of differentiation 11b (CD11b) for monocyte subset characterization. Black box denotes flow cytometry gates used to distinguish $Ly6C^{hi}$ and $Ly6C^{lo}$ monocyte populations.

Supplemental Tables

**Table S1. Three-way and two-way ANOVA analyses of Aβ plaque burden in the prefrontal cortex, hippocampus, and hypothalamus. Related to Figure 1 and S1.**

Three-way ANOVA was used to assess the effects of genotype, sex, and lighting on Aβ plaque burden in the prefrontal cortex, hippocampus, and hypothalamus. Two-way ANOVA results for sex and lighting (and their interaction) are also shown for each region. The table reports F-statistics and corresponding P-values for all main effects and interaction terms.

| 3-way ANOVA | Source of Variation | Prefrontal Cortex | Hippocampus | Hypothalamus |
|---|---|---|---|---|
| Genotype | F-statistic | 77.00 | 211.10 | 2.02 |
| | p-value | <0.001 | <0.001 | 0.16 |
| Sex | F-statistic | 14.43 | 80.44 | 2.02 |
| | p-value | 0.001 | <0.001 | 0.16 |
| Lighting | F-statistic | 6.78 | 42.51 | 2.02 |
| | p-value | 0.013 | <0.001 | 0.16 |
| Genotype x Sex | F-statistic | 14.43 | 80.44 | 2.02 |
| | p-value | 0.001 | <0.001 | 0.16 |
| Genotype x Lighting | F-statistic | 6.78 | 42.51 | 2.02 |
| | p-value | 0.013 | <0.001 | 0.16 |
| Sex x Lighting | F-statistic | 14.82 | 65.19 | 2.02 |
| | p-value | < 0.001 | <0.001 | 0.16 |
| Genotype x Sex x Lighting | F-statistic | 14.82 | 65.19 | 2.02 |
| | p-value | < 0.001 | <0.001 | 0.16 |
| **2-way ANOVA** | **Statistic** | **Prefrontal Cortex** | **Hippocampus** | **Hypothalamus** |
| Interaction | F-statistic | 14.82 | 61.99 | 0.98 |
| | p-value | 0.001 | <0.001 | 0.34 |
| Sex | F-statistic | 14.43 | 76.48 | 0.98 |
| | p-value | 0.001 | <0.001 | 0.34 |
| Lighting | F-statistic | 6.78 | 40.43 | 0.98 |
| | p-value | 0.017 | <0.001 | 0.34 |

**Table S2. Three-way and two-way ANOVA analyses of microglia cell counts in the prefrontal cortex, hippocampus, and hypothalamus. Related to Figure 2.**

Three-way ANOVA was used to assess the effects of genotype, sex, and lighting on microglia cell counts in the prefrontal cortex, hippocampus, and hypothalamus. Two-way ANOVA results for sex and lighting (and their interaction) are also shown for each region. The table reports F-statistics and corresponding P-values for all main effects and interaction terms.

| 3-way ANOVA | Source of Variation | Prefrontal Cortex | Hippocampus | Hypothalamus |
|---|---|---|---|---|
| Genotype | F-statistic | 8.5420 | 1.2960 | 0.1539 |
| | p-value | 0.0057 | 0.2619 | 0.6970 |
| Sex | F-statistic | 1.6790 | 0.1161 | 8.1900 |
| | p-value | 0.2025 | 0.7351 | 0.0067 |
| Lighting | F-statistic | 2.9860 | 0.1075 | <0.0001 |
| | p-value | 0.0917 | 0.7448 | 0.9941 |
| Genotype x Sex | F-statistic | 0.9369 | 0.9818 | 0.3677 |
| | p-value | 0.3389 | 0.3279 | 0.5477 |
| Genotype x Lighting | F-statistic | 2.4490 | 0.4880 | 0.1700 |
| | p-value | 0.1255 | 0.4890 | 0.6823 |
| Sex x Lighting | F-statistic | 4.8500 | 2.7620 | 0.1814 |
| | p-value | 0.0335 | 0.1046 | 0.6724 |
| Genotype x Sex x Lighting | F-statistic | 2.1850 | 0.3003 | 1.1310 |
| | p-value | 0.1472 | 0.5868 | 0.2940 |
| **2-way ANOVA** | **Statistic** | **Prefrontal Cortex** | **Hippocampus** | **Hypothalamus** |
| Interaction | F-statistic | 4.6880 | 3.2020 | 0.9197 |
| | p-value | 0.0426 | 0.0895 | 0.3490 |
| Sex | F-statistic | 1.7740 | 0.2771 | 2.1100 |
| | p-value | 0.1979 | 0.6047 | 0.1619 |
| Lighting | F-statistic | 3.7530 | 0.6908 | 0.0730 |
| | p-value | 0.0670 | 0.4162 | 0.7897 |

**Table S3. Three-way and two-way ANOVA analyses of macrophage cell counts in the prefrontal cortex, hippocampus, and hypothalamus. Related to Figure 2.**

Three-way ANOVA was used to assess the effects of genotype, sex, and lighting on macrophage cell counts in the prefrontal cortex, hippocampus, and hypothalamus. Two-way ANOVA results for sex and lighting (and their interaction) are also shown for each region. The table reports F-statistics and corresponding P-values for all main effects and interaction terms.

| 3-way ANOVA | Source of Variation | Prefrontal Cortex | Hippocampus | Hypothalamus |
|---|---|---|---|---|
| Genotype | F-statistic | 18.01 | 15.22 | 0.04411 |
| | p-value | <0.0001 | 0.0004 | 0.8347 |
| Sex | F-statistic | 10.16 | 2.26 | 3.234 |
| | p-value | 0.0028 | 0.1406 | 0.0797 |
| Lighting | F-statistic | 15 | 21.89 | 2.718 |
| | p-value | 0.0004 | <0.0001 | 0.1071 |
| Genotype x Sex | F-statistic | 7.245 | 2.933 | 0.5699 |
| | p-value | 0.0103 | 0.0945 | 0.4547 |
| Genotype x Lighting | F-statistic | 2.024 | 4.189 | 2.495 |
| | p-value | 0.1625 | 0.0473 | 0.1221 |
| Sex x Lighting | F-statistic | 2.088 | 3.136 | 0.1053 |
| | p-value | 0.1563 | 0.0842 | 0.7473 |
| Genotype x Sex x Lighting | F-statistic | 4.291 | 5.902 | 0.1906 |
| | p-value | 0.0448 | 0.0197 | 0.6647 |
| **2-way ANOVA** | **Statistic** | **Prefrontal Cortex** | **Hippocampus** | **Hypothalamus** |
| Interaction | F-statistic | 4.069 | 5.557 | 0.00631 |
| | p-value | 0.0573 | 0.0287 | 0.9375 |
| Sex | F-statistic | 11.37 | 3.257 | 0.5465 |
| | p-value | 0.003 | 0.0862 | 0.4684 |
| Lighting | F-statistic | 9.229 | 14.24 | 5.229 |
| | p-value | 0.0065 | 0.0012 | 0.0332 |

**Table S4. Summary of ECHO-modeled oscillation parameters for core clock proteins. Related to Figure 3.**

Table reports Extended Circadian Harmonic Oscillator (ECHO) outputs, including oscillation type, fitted circadian periods (hours), and BH-adjusted p-values for rhythmicity for BMAL1, CLOCK, CRY1, CRY2, PER1, and PER3 in male (♂) and female (♀) mice across the LM, AP, CD and CD/AP groups. Dashes denote proteins not detected by tandem mass tag mass spectrometry. Period values of 27 hours represent upper-bound convergence limits of the model. BH-adjusted p-values indicate rhythmicity significance after multiple-testing correction. Fill color indicates genotype/lighting condition (dark blue = LM, light blue = AP, red = CD, orange = CD/AP).

| Protein | Sex | Variable | LM | AP | CD | CD/AP |
|---|---|---|---|---|---|---|
| BMAL1 | ♂ | Oscillation Type | **Forced** | Repressed | Overexpressed | **Damped** |
| | | Period | **18.9603** | 9.4320 | 6.9788 | **27.0000** |
| | | BH Adj p-Value | **0.0040** | 0.0930 | 0.4566 | **0.0001** |
| | ♀ | Oscillation Type | Damped | **Damped** | Damped | **Harmonic** |
| | | Period | 10.2887 | **26.6733** | 11.8358 | **25.3550** |
| | | BH Adj p-Value | 0.3391 | **0.0031** | 0.0643 | **0.0001** |
| CLOCK | ♂ | Oscillation Type | **Forced** | **Harmonic** | Overexpressed | **Harmonic** |
| | | Period | **26.213** | **22.135** | 21.664 | **27** |
| | | BH Adj p-Value | **0.0008** | **0.0205** | 0.0012 | **0.0005** |
| | ♀ | Oscillation Type | - | **Forced** | - | **Harmonic** |
| | | Period | - | **26.4237** | - | **23.0400** |
| | | BH Adj p-Value | - | **0.0001** | - | **0.0001** |
| CRY1 | ♂ | Oscillation Type | - | **Damped** | - | Repressed |
| | | Period | - | **26.2791** | - | 11.1574 |
| | | BH Adj p-Value | - | **0.0001** | - | 0.0039 |
| | ♀ | Oscillation Type | **Forced** | Overexpressed | **Forced** | **Harmonic** |
| | | Period | **22.0573** | 20.9853 | **27.0000** | **26.2060** |
| | | BH Adj p-Value | **0.0001** | 0.0016 | **0.0003** | **0.0001** |
| CRY2 | ♂ | Oscillation Type | **Harmonic** | **Damped** | **Harmonic** | **Damped** |
| | | Period | **19.3465** | **22.6261** | **19.3747** | **27.0000** |
| | | BH Adj p-Value | **0.0001** | **0.0060** | **0.0004** | **0.0001** |
| | ♀ | Oscillation Type | **Harmonic** | - | **Forced** | - |
| | | Period | **22.0830** | - | **18.0225** | - |
| | | BH Adj p-Value | **0.0004** | - | **0.0217** | - |
| PER1 | ♂ | Oscillation Type | Forced | Harmonic | **Damped** | **Forced** |
| | | Period | 11.4249 | 10.8440 | **21.8026** | **27.0000** |
| | | BH Adj p-Value | 0.0060 | 0.0213 | **0.0025** | **0.0002** |
| | ♀ | Oscillation Type | Damped | **Harmonic** | **Damped** | **Harmonic** |
| | | Period | 27.0000 | **24.6875** | **18.9832** | **22.2774** |
| | | BH Adj p-Value | 0.0002 | **0.0030** | **0.0172** | **0.0001** |
| PER3 | ♂ | Oscillation Type | **Harmonic** | - | **Forced** | - |
| | | Period | **21.7714** | - | **23.7034** | - |
| | | BH Adj p-Value | **0.0011** | - | **0.0025** | - |
| | ♀ | Oscillation Type | - | **Damped** | - | **Harmonic** |
| | | Period | - | **27.0000** | - | **27.0000** |
| | | BH Adj p-Value | - | **0.0030** | - | **0.0210** |

**Table S5. Summary of all oscillation and circadian oscillation proteins. Related to Figure 3.**

Table reports the number and percentage of oscillatory proteins from all detected male (♂) and female (♀) LM, AP, CD, and CD/AP proteins before (all oscillations) and after filtering out repressed and overexpression oscillation types (circadian oscillations). Fill color indicates genotype/lighting condition (dark blue = LM, light blue = AP, red = CD, orange = CD/AP).

| Variable | Sex | LM | AP | CD | CD/AP |
|---|---|---|---|---|---|
| **All Oscillations** *(Harmonic/Damped/Forced/ Repressed/Overexpressed)* | ♂ | 2038 | 1883 | 1796 | 2406 |
| | ♀ | 1791 | 1822 | 1780 | 2171 |
| **Circadian Oscillations** *(Harmonic/Damped/Forced)* | ♂ | 1877 | 1764 | 1545 | 2077 |
| | ♀ | 1707 | 1670 | 1610 | 2052 |
| **All Oscillations (%)** | ♂ | 39.68 | 36.66 | 34.97 | 46.85 |
| | ♀ | 34.87 | 35.48 | 34.66 | 42.27 |
| **Circadian Oscillations (%)** | ♂ | 36.53 | 34.35 | 30.08 | 40.44 |
| | ♀ | 33.24 | 32.52 | 31.35 | 39.95 |
| **Difference (%)** | ♂ | 3.13 | 2.32 | 4.89 | 6.41 |
| | ♀ | 1.64 | 2.96 | 3.31 | 2.32 |

**Table S6. Pairwise comparisons of circadian phase shifts between lighting groups. Related to Figure 4.**

Kruskal–Wallis tests indicated significant differences in ΔCT-based phase shifts across the four experimental pairings (male (♂) LM vs CD, male AP vs CD/AP, female (♀) LM vs CD, and female AP vs CD/AP). Post-hoc pairwise comparisons were performed using Dunn's test with FDR correction, and the table shows the FDR-adjusted p-values for each comparison. Color indicates lighting comparison (purple = ♂ LM vs CD, green = ♂ AP vs CD/AP, pink = ♀ LM vs CD, orange = ♀ AP vs CD/AP).

| Contrast | ♂ LM vs CD | ♂ AP vs CD/AP | ♀ LM vs CD | ♀ AP vs CD/AP |
|---|---|---|---|---|
| ♂ LM vs CD | 1.0000 | 0.2103 | 0.6445 | 0.0138 |
| ♂ AP vs CD/AP | 0.2103 | 1.0000 | 0.3978 | 0.0001 |
| ♀ LM vs CD | 0.6445 | 0.3978 | 1.0000 | 0.0047 |
| ♀ AP vs CD/AP | 0.0137 | 0.0001 | 0.0047 | 1.0000 |

**Table S7. Circadian oscillation parameters for phagocytosis, mitochondrial membrane potential (ΔΨm), and reactive oxygen species (ROS). Related to Figure 6.**

Circadian oscillations were measured in male (♂) and female (♀) mice across four experimental groups: LM, AP, CD, and CD/AP. For each variable, oscillation type indicates the classification of the rhythmic pattern (harmonic, repressed, forced, damped, or overexpressed). Period represents the duration of one oscillation cycle in hours, hours shifted indicates the phase shift relative to baseline in hours, and adj p-value is the Benjamini-Hochberg adjusted p-value for oscillation significance. Each row corresponds to a specific sex and experimental group for the indicated variable. Fill color indicates genotype/lighting condition (dark blue = LM, light blue = AP, red = CD, orange = CD/AP).

| Experiment | Sex | Variable | LM | AP | CD | CD/AP |
|---|---|---|---|---|---|---|
| Phagocytosis | ♂ | Oscillation Type | **Harmonic** | Repressed | Harmonic | Harmonic |
| | | Period | **30.0000** | 18.6509 | 22.2570 | 21.7089 |
| | | Hours Shifted | **12.1197** | 10.5014 | 13.1275 | 12.4878 |
| | | BH Adj p-Value | **0.0062** | 0.4527 | 0.3175 | 0.2401 |
| | ♀ | Oscillation Type | Harmonic | Harmonic | Harmonic | Forced |
| | | Period | 22.3192 | 24.4178 | 29.9541 | 15.6000 |
| | | Hours Shifted | 4.6926 | 7.6154 | 12.3577 | 5.3753 |
| | | BH Adj p-Value | 0.1030 | 0.3301 | 0.5395 | 0.5133 |
| ΔΨm | ♂ | Oscillation Type | **Harmonic** | Repressed | **Harmonic** | **Harmonic** |
| | | Period | **25.0445** | 12.0000 | **23.7423** | **26.5923** |
| | | Hours Shifted | **5.4591** | 9.9433 | **11.8608** | **15.0798** |
| | | BH Adj p-Value | **0.0031** | 0.5827 | **0.0130** | **0.0009** |
| | ♀ | Oscillation Type | **Forced** | **Overexpressed** | **Damped** | **Damped** |
| | | Period | **30.0000** | **30.0000** | **18.9190** | **21.4347** |
| | | Hours Shifted | **1.4714** | **1.9377** | **10.2868** | **8.8186** |
| | | BH Adj p-Value | **0.0048** | **0.0073** | **0.0301** | **0.0015** |
| ROS | ♂ | Oscillation Type | **Forced** | **Overexpressed** | **Harmonic** | **Harmonic** |
| | | Period | **21.0407** | **28.5169** | **24.6946** | **24.6938** |
| | | Hours Shifted | **14.6634** | **4.2812** | **14.4938** | **14.9359** |
| | | BH Adj p-Value | **0.0002** | **0.0097** | **0.0002** | **0.0002** |
| | ♀ | Oscillation Type | **Harmonic** | **Harmonic** | **Forced** | **Forced** |
| | | Period | **25.2460** | **25.4034** | **25.5839** | **28.7798** |
| | | Hours Shifted | **4.4029** | **4.3766** | **6.1730** | **1.9265** |
| | | BH Adj p-Value | **0.0002** | **0.0002** | **0.0013** | **0.0004** |

**Table S8. Statistical analysis of male daily mean reactive oxygen species (ROS) values. Related to Figure 6.**

Three-way ANOVA was performed to evaluate the effects of sex, genotype, and lighting on daily average ROS. Subsequent two-way ANOVAs were conducted separately for females (♀) and males (♂) to examine genotype × lighting interactions. The table reports F-statistics and corresponding p-values for each main effect and interaction.

| 3-way ANOVA | F stat | p value |
|---|---|---|
| Sex | 2.8050 | 0.1134 |
| Genotype | 0.5865 | 0.4549 |
| Lighting | 0.7666 | 0.3942 |
| Sex x Genotype | 3.0580 | 0.0995 |
| Sex x Lighting | 12.5800 | 0.0027 |
| Genotype x Lighting | 18.5600 | 0.0005 |
| Sex x Genotype x Lighting | 35.7400 | <0.0001 |
| ♀ 2-way Anova | F stat | p value |
| Interaction | 1.3520 | 0.2784 |
| Genotype | 0.4685 | 0.5130 |
| Lighting | 9.4870 | 0.0151 |
| ♂ 2-way Anova | F stat | p value |
| Interaction | 54.6100 | <0.0001 |
| Genotype | 3.2630 | 0.1085 |
| Lighting | 3.6840 | 0.0912 |

**Table S9. Two-way ANOVA of female peripheral monocyte populations. Related to Figure 6.**

Analysis was performed to assess the effects of genotype and lighting on female (♀) non-inflammatory and pro-inflammatory circulating blood monocytes. The table reports F-statistics and corresponding p-values for the interaction term, as well as the main effects of genotype and lighting.

| 2-way ANOVA | Statistic | Non-Inflammatory | Inflammatory |
|---|---|---|---|
| Interaction | F-statistic | 0.2713 | 0.8954 |
| | p-value | 0.6079 | 0.3543 |
| Genotype | F-statistic | 1.1020 | 3.6800 |
| | p-value | 0.3057 | 0.0682 |
| Lighting | F-statistic | 0.4598 | 0.8697 |
| | p-value | 0.5051 | 0.3612 |
